# Enhancers predominantly regulate gene expression *in vivo* via transcription initiation

**DOI:** 10.1101/844191

**Authors:** Martin S. C. Larke, Takayuki Nojima, Jelena Telenius, Jacqueline A. Sharpe, Jacqueline A. Sloane-Stanley, Sue Butler, Robert A. Beagrie, Damien J. Downes, Ron Schwessinger, A. Marieke Oudelaar, Julia Truch, Bryony Crompton, M. A. Bender, Nicholas J. Proudfoot, Douglas R. Higgs, Jim R. Hughes

## Abstract

Gene transcription occurs via a cycle of linked events including initiation, promoter proximal pausing and elongation of RNA polymerase II (Pol II). A key question is how do transcriptional enhancers influence these events to control gene expression? Here we have used a new approach to quantify transcriptional initiation and pausing *in vivo*, while simultaneously identifying transcription start sites (TSSs) and pause-sites (TPSs) from single RNA molecules. When analyzed in parallel with nascent RNA-seq, these data show that differential gene expression is achieved predominantly via changes in transcription initiation rather than Pol II pausing. Using genetically engineered mouse models deleted for specific enhancers we show that these elements control gene expression via Pol II recruitment and/or initiation rather than via promoter proximal pause release. Together, our data show that enhancers, in general, control gene expression predominantly by Pol II recruitment and initiation rather than via pausing.

## INTRODUCTION

Changes in RNA expression during cell fate decisions are made via the activation of promoters by enhancers that respond to changes in transcriptional and epigenetic programmes to regulate gene expression in time and space. Many enhancers physically contact target promoters via changes in the three dimensional conformation of the genome (Hay et al. 2016; Oudelaar et al. 2018; Pennacchio et al. 2013; Rowley and Corces 2018; Shlyueva, Stampfel, and Stark 2014). This may increase the concentration of transcription factors, co-factors, and Pol II at the promoters of their target genes Cho et al. 2018; Cramer 2019; Heinz et al. 2015; J. Li et al. 2019; Ptashne and Gann 1997; Sabari et al. 2018). However, there are many potential points within the transcription cycle at which such high concentrations of transcription factors and co-factors could influence expression including regulating initiation, pausing or elongation (Boehning et al. 2018; Cho et al. 2018; Cisse et al. 2013; Hnisz et al. 2017; Lu et al. 2018; Vernimmen et al. 2007). Recent studies have suggested that one major point of regulation involves releasing pre-loaded, paused Pol II complexes into productive elongation (Cho et al. 2018; Cramer 2019; Heinz et al. 2015; J. Li et al. 2019; Ptashne and Gann 1997; Sabari et al. 2018).

Regulation of the transcription cycle has been a topic of intense research and has been characterized in detail (Chen, Smith, and Shilatifard 2018; Jonkers and Lis 2015; Shandilya and Roberts 2012; Wissink et al. 2019). General transcription factors bind to a nucleosome-free promoter causing conformational changes in the DNA helix that allow the binding of Pol II and concomitant synthesis of nascent RNA from the TSS. This pre initiation complex (PIC) is modified by phosphorylation and transitions from an initiating complex into an elongating complex (Buratowski et al. 1989; Sainsbury, Bernecky, and Cramer 2015). The transition from initiation to elongation is thought to be an important point of control of gene expression (Chen, Smith, and Shilatifard 2018; Fraser, Sehgal, and Darnell 1978; Gariglio, Bellard, and Chambon 1981; Jonkers and Lis 2015). One potential point of regulation occurs +30-60 (bps) downstream of the TSS where the nascent elongation complex, including Pol II and a short 7-Methylguanosine (m^7^G) capped RNA molecule, transiently pauses via the action of negative elongation factors (NELF and DSIF) (Yamaguchi et al. 1999; T. Wada et al. 1998). The paused Pol II complex is then converted into an elongation competent form through phosphorylation of its constituent factors by CDK9, the kinase component of the positive transcription elongation factor (P-TEFb). Transcriptional pausing is thereby released (Marshall and Price 1995) allowing productive elongation (Adelman and Lis 2012; Chen, Smith, and Shilatifard 2018; Jonkers and Lis 2015). The key question addressed here is how do enhancers control this transcription cycle at their cognate promoters?

Current evidence suggests that pause-release may be a major point of enhancer mediated regulation. Genome-wide ChIP-seq analysis has shown that Pol II accumulates across the promoter proximal region (∼-100-+300) including the TSSs and TPSs of most protein coding genes (Chen, Smith, and Shilatifard 2018; L. Core and Adelman 2019; Fraser, Sehgal, and Darnell 1978; Gariglio, Bellard, and Chambon 1981; Jonkers and Lis 2015; Kim et al. 2005; Muse et al. 2007; Wissink et al. 2019; Zeitlinger et al. 2007). It has been proposed that the density of Pol II found at the promoter proximal region represents paused Pol II and the ratio of this to the density of Pol II throughout the remainder of the body of the gene (the pausing index) has been used as a measure of transcriptional pausing (Muse et al. 2007; Rahl et al. 2010; Zeitlinger et al. 2007). Inhibition of CDK9 causes the pausing index to increase, and this has been interpreted as evidence for regulation of transcriptional pausing (Gressel et al. 2017; Jonkers, Kwak, and Lis 2014; Rahl et al. 2010). Consequently, changes in the pausing index have been used to suggest that pausing is a key regulatory step in the transcription cycle (Adelman and Lis 2012; Chen, Smith, and Shilatifard 2018; Jonkers and Lis 2015) and that this is regulated by enhancers (Chen et al. 2017; Gao et al. 2018; Ghavi-Helm et al. 2014; Liu et al. 2013; Schaukowitch et al. 2014). Clearly this interpretation depends on whether the promoter-proximal peak of Pol II observed in ChIP experiments reflects initiation, pausing, or both (Ehrensberger, Kelly, and Svejstrup 2013). Considering that pausing occurs only 30-60nts downstream of initiation, the resolution provided by Pol II ChIP-seq is not best suited to resolve these two events in the transcription cycle (Wissink et al. 2019). Furthermore, the level of Pol II bound does not necessarily reflect the level of transcription. Therefore, our ability to measure the levels of RNA at each of these steps of the transcription cycle is critical to understand how enhancers regulate the activation of gene expression.

Here, using a newly developed genome-wide assay that measures short capped RNAs (scaRNA-seq), we accurately and simultaneously identify TSSs and TPSs from single molecules at high resolution. By comparing scaRNA-seq data with measurements of nascent transcription using mNET-seq (Nojima et al. 2015) and intronic RNA (Boswell et al. 2017; La Manno et al. 2018), we find that enhancer-driven transcription, during in vivo differentiation of primary cells, appears to be controlled predominantly via initiation rather than pause-release or elongation. To test this experimentally we investigated the effect of the loss the major enhancers of the well-characterized globin loci on transcriptional initiation and pausing in primary erythroid cells. Previous work has identified the major enhancers of the α- and β-globin enhancer clusters, whose deletion results in reduction of accumulated and nascent transcription (Bender et al. 2012; Hay et al. 2016). We compared the levels of scaRNA and nascent RNA in mouse models, in which either the major α- or β-globin enhancers have been deleted with wild type animals. Consistent with the conclusions from genome-wide studies, we show that in each case, the enhancers primarily affect the stages of recruitment and/or initiation of transcription rather than pause-release or elongation.

## RESULTS

### Analysis of transcription initiation and Pol II pausing *in vivo* using scaRNA-seq

To understand how initiation and promoter proximal pausing affect gene expression, it is necessary to measure these distinct stages of transcription quantitatively, at high resolution, and at single genes *in vivo.* To date, assays such as Start-seq (Henriques et al. 2013; Nechaev et al. 2010) can measure initiation and pausing, very close to the TSS (<=+100bps) by sequencing RNA 5’ and 3’ ends separately. To examine initiation and pausing simultaneously and across a larger window, we modified the Start-seq assay. First, we isolated much larger RNA transcripts (<=300nt compared to ∼100nt), enriching for nascent RNA and enabling us to identify the distribution of RNA 3’ ends far beyond the predicted range of Pol II pausing (+30-60bps relative to the TSS), and also beyond the boundary of the +1 nucleosome (+ ∼150bps) (Jonkers and Lis 2015; Mieczkowski et al. 2016; Voong et al. 2016; Yazdi et al. 2015). Second, we used paired-end sequencing to identify both ends of each nascent RNA transcript capturing individual episodes of transcriptional initiation and pausing from single molecules. We refer to this modified assay as “short capped RNA sequencing” (scaRNA-seq; Figure 1A).

**Figure 1.**
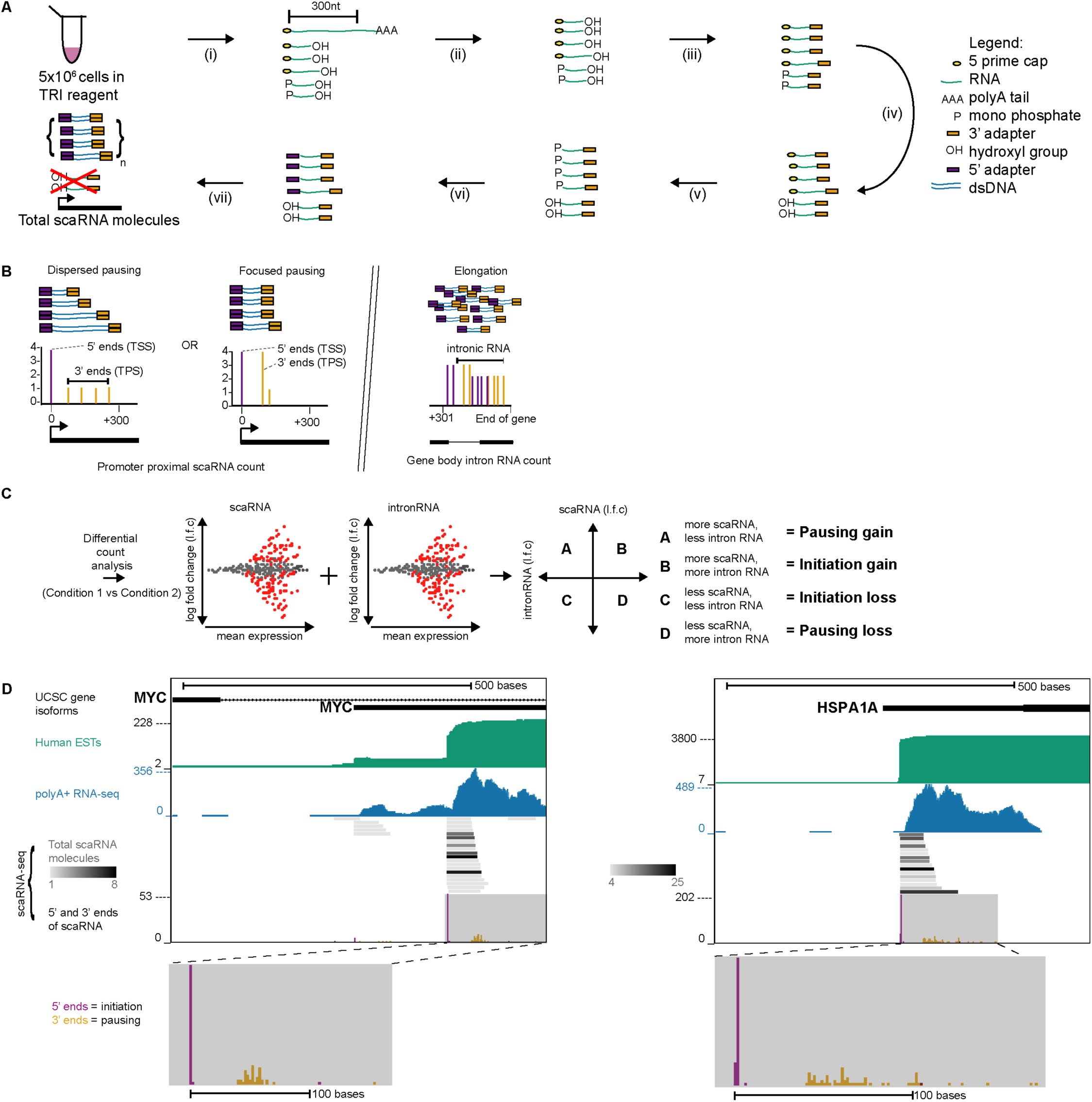
scaRNA-seq maps transcription initiation and pausing at a single molecule level *in vivo*. (A) A schematic of the scaRNA-seq procedure showing key stages. i) 5×10^6^ cells are lysed in TRI reagent and total RNA is extracted. ii) Transcripts <=300nts are selected using denaturing polyacrylamide gel electrophoresis, enriching for nascent RNAs. iii) An Illumina sequencing adapter is ligated onto a free hydroxyl group, marking the 3’ end of the RNA molecule. iv) Phosphatase treatment prevents degraded or abundant RNA such as ribosomal RNA from entering downstream adapter ligation step. v) The m^7^G cap is converted into a monophosphate using the CAP-clip enzyme. vi) An Illumina sequencing adapter is ligated to 5’ monophosphates. vii) Adapter ligated scaRNAs are reverse transcribed into cDNA and amplified using PCR to generate libraries for sequencing. Alignment of read pairs genome wide allows mapping and counting of scaRNA molecules. (B) Measuring promoter proximal and distal transcription: Localization of initiation and pausing is achieved using the extreme 5’ and 3’ bases of each reconstructed scaRNA molecule. Pausing may occur in dispersed or focused modes. Total counts of scaRNA molecules in promoter proximal regions (0-300bp, relative to observed TSS) stand proxy for levels of initiation and pausing. intronic RNA is used as a measure of elongation. (C) Integrating transcriptional data: scaRNA and intronicRNA counts (expression) were analyzed using scaRNA derived (overserved) TSS. Differential scaRNA and nascent RNA counts analysis were performed comparing datasets in condition when gene expression changes; in this case 0h and 24h of erythropoiesis. Significant changes in expression are represented as red dots. Comparing each scaRNA log fold change to the associated log fold change in intronicRNA are shown as a scatter plot. Each quadrant identifies a distinct class of regulation; A (pausing gain), B (Initiation gain), C (Initiation loss), D (pausing loss). (D) scaRNA-seq in K562 cells. Left hand side shows MYC and right-hand side shows HSPA1A (also known as Hsp70). The figure shows annotated UCSC gene isoforms, Human expressed sequence tags (ESTs), polyA+ RNA-seq in K562 cells, the density of reconstructed RNA molecules and distributions of RNA 5’ and 3’ ends, indicative of punctate initiation and promoter proximal pausing at expected positions downstream of the site of initiation (+20-60nt). PolyA+ data Y axis = number mapped reads (raw data). scaRNA data Y axis = Number of RNA 5’ or 3’ ends (raw data).

As set out in the model (Figure 1A), using this assay, it is possible to count total numbers of individual molecules as a simple measure of promoter proximal transcription. However, the 5’ or 3’ ends of these molecules can also be visualized as separate distributions, yielding maps of TSSs and TPSs respectively. If there were no pausing, the 3’ ends of initiating transcripts would be evenly distributed across the entire ∼300nt promoter proximal window due to an even rate of progression. When transcription pauses this would be observed as a peak, or peaks of clustered 3’ ends within the promoter proximal region (Figure 1B).

Changes in either initiation or pausing rates will affect the number of promoter proximal transcripts observed by scaRNA-seq. The overall level of promoter proximal transcription (initiated and paused RNA) can then be directly compared with the associated levels of intronic RNA counts, which act as a measure of nascent transcription. Gene regulation due to changes in initiation would show a positive correlation between the levels of scaRNAs and intronic RNA, in that increases in promoter proximal scaRNA transcripts would lead to increases in overall gene expression or *vice-versa*. However, gene regulation due to changes in promoter proximal pausing would show as a negative correlation between the levels of scaRNAs and the overall output of the gene (Figure 1C) as paused transcripts build up at the expense of elongation and full-length length transcription.

### Validating scaRNA-seq in genome-wide assays

We validated this assay by analysing K562 cells since there are many previous datasets analysing TSSs, initiation, and pausing in this cell line (L. J. Core et al. 2014; Tome, Tippens, and Lis 2018). We first looked at genes whose initiation and pausing had been previously characterized in K562 cells: in particular, using scaRNA-seq we examined the *HSPA1A* (Heat shock protein 70) and *MYC* genes (Figure 1D). At each gene this showed a predominant punctate TSS (purple peaks) and the linked downstream TPSs (orange peaks) consistent with Pol II pausing. We thus confirmed that scaRNA-seq can detect and quantify transcription initiation and Pol II pausing simultaneously at individual genes and at base pair resolution using a modest number of cells (5×10^6^).

To validate the ability of scaRNA-seq to accurately detect transcription start sites we called TSSs from scaRNA 5’ ends genome-wide using Homer (Heinz et al. 2010) and selected those that lie within 500bp of previously annotated UCSC gene TSSs. We then scored each called TSS for the presence of Initiator and TATA box motifs (Vo Ngoc et al. 2017) and observed a strong enrichment at the TSSs called from scaRNA-seq data (Figure S1A). Next, to assess the ability of scaRNA-seq to detect transcription pause sites in K562, we generated mNET-seq which identifies 3’ RNA ends associated with Pol II (Nojima et al. 2015). We also analyzed previously published GRO-seq data, which maps nascent RNA 3’ ends using an *in vitro* run on assay (L. J. Core et al. 2014; L. J. Core, Waterfall, and Lis 2008). The distribution of scaRNA 3’ ends fully corresponds to the profiles obtained using these previously validated approaches (Figure S1B) confirming that scaRNA-seq can simultaneously and accurately map both TSSs and TPSs *in vivo*.

### Use of scaRNA-seq revises the model of promoter proximal transcription

Visualization of *HSPA1A* and *MYC* genes revealed an interesting phenomenon; at both genes, the predominant RNA 5’ end signal (observed using scaRNA-seq) is located on the 5’ side of the TSSs annotated in UCSC; supported by both analysis of human ESTs and polyA+ RNA-seq. The UCSC annotated *MYC* TSS appears to be a minor TSS in K562 cells confirmed by scaRNAs, ESTs and polyA+ RNA-seq. For the *HSPA1A* gene the TSS is poorly supported by any of the annotations. We then plotted the relative positions of all TSSs observed in K562 cells using scaRNA-seq versus those annotated in UCSC. This showed a striking systematic skew with more than 94% of TSS observed with scaRNA-seq lying up or downstream of UCSC annotated TSSs: less than 6% (660/11165) of the UCSC annotated TSSs map to the precise base pair observed using scaRNA-seq (Figure 2A). However, the TSSs called from scaRNA-seq data had a 3-4x higher enrichment for the TSS associated Initiator and TATA motifs than annotated UCSC TSSs (Figure S1A). This suggests that scaRNA-seq provides a better way of accurately finding the most frequently used TSSs than using these general database gene annotations which are commonly used in the meta-analysis of genomics data (Adelman and Lis 2012; L. J. Core et al. 2014; Jonkers and Lis 2015; Rahl et al. 2010). We therefore investigated whether using observed (scaRNA-seq) rather than annotated TSSs would alter our interpretation of promoter proximal transcription.

**Figure 2.**
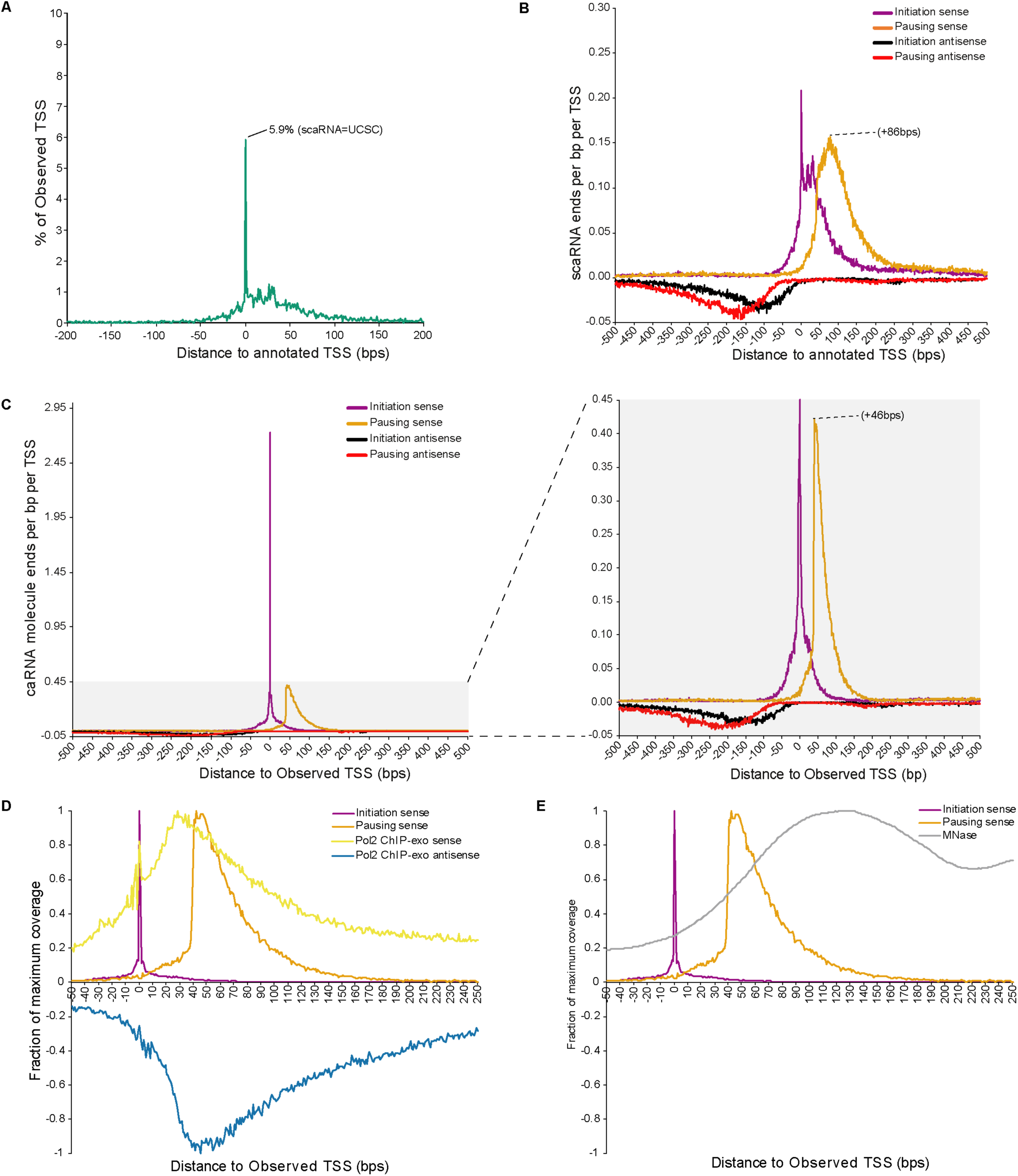
Annotated TSS positions are systematically skewed upstream of their *in vivo* locations. (A) A plot of the distance between the position of observed TSSs (called from scaRNA-seq data) lying within (+/-500bps) of the previously annotated TSSs associated with UCSC genes (n=11165). Only 5.9% of TSS overlap with previously annotated UCSC TSSs, suggesting a systematic skew of annotated TSSs mostly upstream of the observed TSSs. (B) A meta-analysis of scaRNA-seq data around the annotated UCSC TSSs for which an observed TSS could be found (n = 11165). This analysis shows dispersed initiation around the TSS (RNA 5’ ends in purple), promoter proximal accumulation of RNA 3’ ends (Pol II pausing) at variable distances from the TSS (20-200nt) and a maximum signal at +86bps, and high levels of antisense transcription (RNA 5’ (black) and 3’ ends (red)). Data displayed as raw (unnormalized) coverage of RNA ends (reads per bp per TSS). (C) A meta-analysis of scaRNA-seq data around observed TSS positions (n = 11165), showing punctate initiation around the TSS and a loss of diffuse RNA 5’ ends signal downstream of the TSS after correcting for discrepancies in the positions of TSSs. Pol II pausing predominantly occurs at +46bps and extends to around +150bps relative to the TSS. Antisense transcription is still seen but is much less as a proportion of the sense transcription. (D) A meta-analysis of scaRNA-seq RNA 5’ and 3’ ends on the sense strand. The data are plotted relative to observed (scaRNA-seq) TSS. The Y-axis is scaled to 1 for 5’ or 3’ ends to simplify visualization. ChIP-exo data for Pol II, (sense strand reads are shown in yellow, antisense reads in blue). The regions of maximum Pol II occupancy lie between the maximum signals on the sense strand and minimum signals on antisense strand; 0bp (TSS Pol II) and ∼+35-55bp (Paused Pol II). ChIP-exo data are re-analyses of previously published data (Mchaourab, Perreault, and Venters 2018; Rhee and Pugh 2012). (E) A meta-analysis of scaRNA-seq RNA 5’ and 3’ ends on the sense strand. The data are plotted relative to observed (scaRNA-seq) TSS. The Y-axis is scaled to 1 for 5’ or 3’ ends to simplify visualization. MNase-seq data highlighting nucleosome occupancy is represented by a grey line. Maximum MNase coverage is observed around 130bo relative to the TSS, coinciding with the scaRNA 3’ RNA end distribution dropping to baseline. MNase data are re-analyses of previously published data (Mieczkowski et al. 2016).

Plotting scaRNA-seq data with reference to UCSC annotated TSSs we recapitulated the generally accepted view of transcription. UCSC TSSs showed dispersed transcription initiation around each TSS (RNA 5’ ends), and similar dispersed pausing +0-50bps downstream of TSSs (3’ RNA ends; Figure 2B and S1B). Strikingly, scaRNA-seq (observed) TSSs give significantly different meta profiles (Figure 2C). First, transcription initiation (5’ RNA ends) on the sense strand appears to be extremely focused rather than dispersed around the TSS with most of the 5’ ends localized to within a few base pairs of the predominant observed TSS (Figure 2C and Figure S2B). To investigate this effect, we classified TSSs as focused if they contained >=50% of the RNA 5’ ends within a +/-2nt window around the TSS or dispersed if <50% of RNA 5’ ends fell within the +/-2nt window (Figure S2A). As discussed above, when using UCSC annotated TSSs for this analysis the predominant mode of transcription initiation appears to be dispersed. However, using TSSs observed with scaRNA-seq there is an approximately equal division between focused and dispersed TSSs, consistent with evidence suggesting genes may initiate in either or both modes in approximately equal proportions (L. J. Core et al. 2014; Haberle and Stark 2018). Visualization of data from other TSS enrichment methods in this cell type (CAGE and GRO-cap) over annotated UCSC and scaRNA-seq TSSs (S1C and S1D) confirms that scaRNA-seq derived TSSs change our interpretation of the nature of transcription initiation, regardless of the assay used to map TSSs. This implies that a systematic bias has arisen during the pruning of canditate gene TSS positions in the UCSC annotations.

Furthermore, using observed (scaRNA-seq) rather than UCSC annotated TSSs showed that RNA 3’ ends consistently accumulate at +46nt downstream of the TSS (indicative of Pol II pausing) and then decay towards +150nt downstream of the TSS (Figure 2C and S2B). This suggests that Pol II pausing rarely occurs before +46nt relative to the TSS and terminates within ∼100nt of this position. To examine this further, we reanalyzed Pol II ChIP-exo data, which provides a high resolution map of protein binding within chromatin (Figure 2D and S2C; Rhee and Pugh 2011). This demonstrated that Pol II occupancy coincides with the maximum RNA 3’ end signal at +46nt, confirming that Pol II commonly accumulates at this position. An additional peak of Pol II occupancy is also visible at the TSS (Figure 2D) which was previously undetectable using UCSC annotated TSSs (Figure S2D).

To investigate whether the systematic skew in UCSC annotated TSSs was common to other widely used annotations we plotted our scaRNA-seq data over these alternative annotations revealing the same inherent skew in meta profiles (Figure S2E). The severity of the skew depended on the particular annotation being tested. Ensembl TSSs were the least affected and Refseq TSSs the most. This suggests that systematic skew in TSS annotation is present in many widely used annotations and has likely affected our interpretation of promoter proximal transcription due to a loss of resolution in previously published meta profiles.

### Transcriptional pausing at the +1 nucleosome

The distribution of RNA 3’ ends towards the +150nt position prompted us to investigate the level of nucleosome occupancy over the promoter proximal region because the +1 nucleosome has been proposed to play a role in regulation of promoter proximal transcription (Jimeno-González, Ceballos-Chávez, and Reyes 2015; Mieczkowski et al. 2016; Voong et al. 2016; Weber, Ramachandran, and Henikoff 2014). We determined nucleosome density in K562 around observed TSSs using published data (Mieczkowski et al. 2016) and plotted this as a meta profile (Figure 2E) and heatmap (Figure S2B). In both cases this revealed an excellent fit between decreasing density of 3’ scaRNA-seq ends and increasing nucleosome occupancy, which peaks around 120-140bps downstream of the TSS. These observations suggest a model whereby Pol II molecules initiate transcription at a TSS where they experience an elongation bottleneck manifested as Pol II pausing; the extent of Pol II occupancy is limited by the +1 nucleosome with the elongating complex accumulating against the +1 nucleosome. This is consistent with observations suggesting that the +1 nucleosome may enforce Pol II pausing (Jimeno-González, Ceballos-Chávez, and Reyes 2015; Studitsky et al. 2016; Weber, Ramachandran, and Henikoff 2014).

Together, these observations suggest that initiation occurs within a very narrow window around the TSS. Furthermore, elongating transcripts accumulate at +46nt relative to the TSSs suggesting this is the position of Pol II pausing at the majority of promoters. This in turn may be the result of a bottleneck delimited by the +1 nucleosome between ∼+50-150bps relative to the TSS. This would be consistent with previous experiments and computational models of Pol II ChIP-seq data which propose that initiation and promoter proximal pausing are intrinsically linked and pausing may only become rate limiting after initiation is saturated (Ehrensberger, Kelly, and Svejstrup 2013; Gressel et al. 2017; Gressel, Schwalb, and Cramer 2019).

### Differential gene expression is predominantly regulated via recruitment or initiation of transcription rather than Pol II pausing

It has been suggested that regulated Pol II pausing is a ubiquitous mechanism to control gene expression which relies on a stably paused Pol II complex which can be acted on to cause pause release (Adelman and Lis 2012; Jonkers, Kwak, and Lis 2014; Jonkers and Lis 2015; Rahl et al. 2010). However, recently it has also been suggested that the high density of Pol II as revealed by ChIP-seq may reflect a dynamic process of premature termination (Erickson et al. 2018). This is supported by *in vivo* imaging studies which reveal the majority of Pol II is associated with chromatin for seconds (Steurer et al. 2018) and modeling which revealed that the turnover of Pol II on housekeeping genes was comparable to highly “paused” genes such as HSPA1 (Krebs et al. 2017). Whilst it is clear from many studies that Pol II is often found near the TSS the question remains to what extent does this accumulation contribute to changes in gene expression?

To investigate how changes in the level of Pol II pausing contribute to changes in gene expression we used a mouse primary erythroid cell differentiation system (Hay et al. 2016). We generated polyA+ RNA-seq and scaRNA-seq data sets at 0 hour and 24 hour differentiation time points, representing “early” and “late” erythropoiesis (Edling and Hallberg 2007; Hay et al. 2016; McGrath, Catherman, and Palis 2017), to allow for a robust comparison of changes in nascent gene expression and promoter proximal transcription. Nascent RNA was quantified by measuring intronic RNA (Figure S3A, B and C) derived polyA+ RNA sequencing (Boswell et al. 2017; La Manno et al. 2018).

To examine how changes in scaRNA related to changes in nascent transcription at the 0h and 24h timepoints. We performed differential count analysis of intronic RNA for all genes which could be assigned to a scaRNA-seq TSS (Anders and Huber 2010). We next plotted the log fold change (logFC) in scaRNA against the logFC in intron RNA (Figure 3A). This revealed a positive correlation between changes in scaRNA and nascent (intronic) RNA, consistent with a change in initiation at >90% of significantly differentially expressed genes (Figure 1C). For example, both *Npm1* and *Tfrc* have characteristic accumulations of RNA 3’ ends (TPSs) downstream of their TSSs consistent with some degree of Pol II pausing. However, the distribution of 3’ RNA ends (TPSs) does not greatly alter with changes in gene expression (Figure 3B, C). Importantly, the total level of scaRNA changes concordantly with intronic RNA at these genes suggesting regulation through initiation (Figure 1C). A relatively small proportion (<10%) of genes have opposing changes in scaRNA and intronic RNA levels that would be consistent with “Pol II pausing”. However, negative correlations between scaRNA and intronic RNA may still not be due to pausing in this analysis. For example, *Tfdp2* was suggested in this analysis to be upregulated via pause-release (loss of scaRNA, gain in intron RNA) however inspection of the TSS of this genes indicates a loss of scaRNA-seq signal over one TSS, but a gain at a closely spaced upstream TSS, consistent with the use of an alternative promoter rather than Pol II pause-release (Figure S3E). These data suggest that although paused Pol II can be detected at most genes it does not significantly regulate expression of full-length RNA.

**Figure 3.**
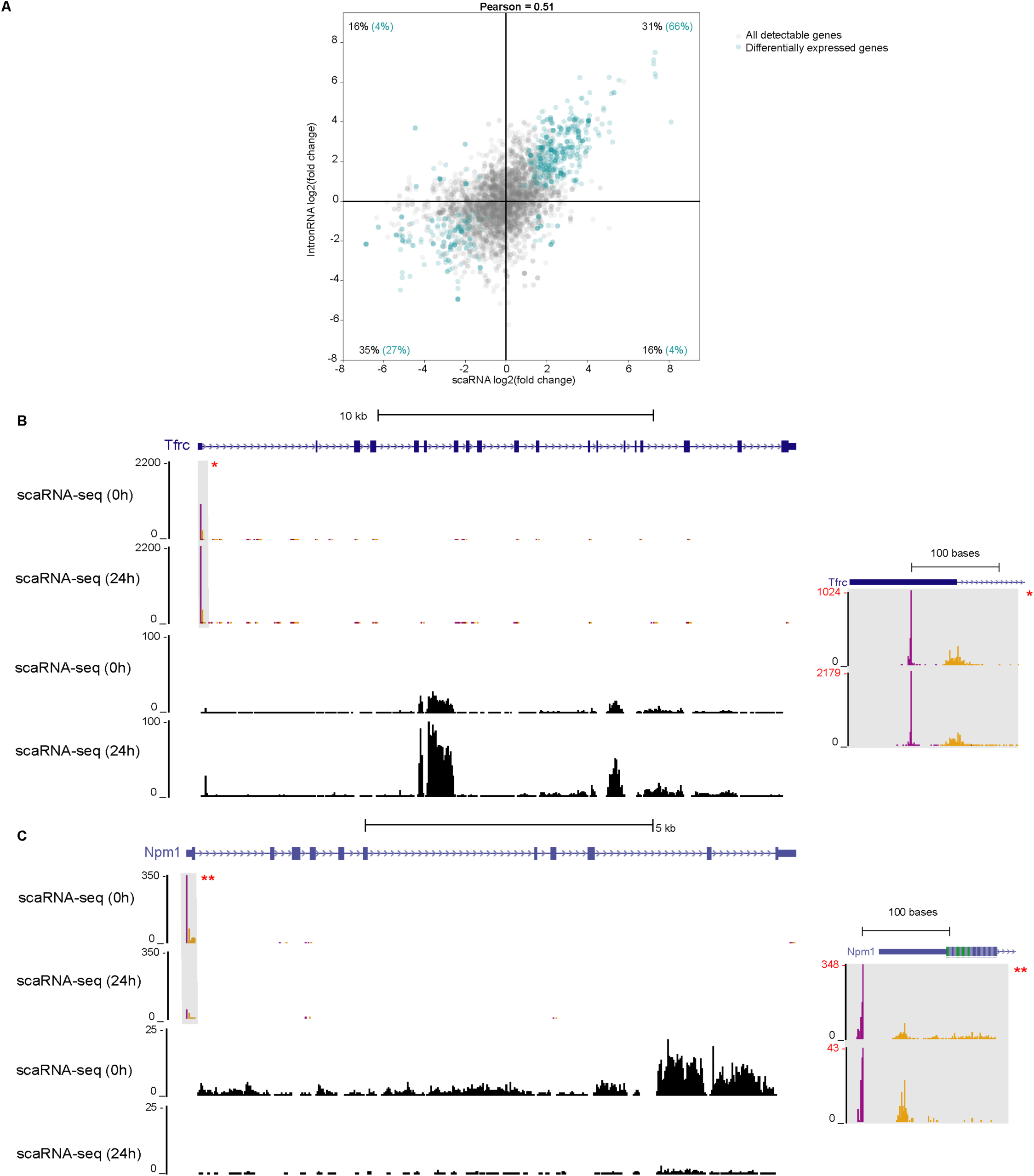
Transcription initiation is the predominant point of regulation in the transcription cycle. (A) A scatter plot of log fold change in scaRNA molecules versus intronic RNA for all detectably expressed genes (grey dots, n=4502). Genes which show significantly differential expression (cyan dots, n=603, p=<0.05 are shown as cyan dots. The data show a positive correlation (Pearson r^2^ of 0.51) between changes in scaRNA and intronic RNA levels. 93% (27%+66%) of differentially expressed genes are positively correlated. As set out in Figure 1B, this shows that regulation via initiation is the predominant step in achieving differential gene expression *in vivo.* (B) *Tfrc* exemplifies increases in scaRNA across the region of promoter proximal transcription and intronic RNA from the remainder of the gene, between 0h and 24h. This indicates an increase in initiation rather than pausing (see Figure 1B). Set inset right for a zoomed view of scaRNA-seq data over promoter (grey box), with Y-axis scaled for visualization. (C) *Npm1* exemplifies decreases in intronic RNA and scaRNA at 0h and 24 h across the region of promoter proximal transcription. The level of scaRNA (5’ and 3’ end) transcripts decrease in parallel with intronic RNA at 24h indicating a decrease in initiation rather than increase in pausing which would be associated with an increase in scaRNA (see Figure 1B). Set inset right for a zoomed view of scaRNA-seq data over promoter (grey box), with Y-axis scaled for visualization. All data were derived from biological triplicates at each stage (0h or 24h) of erythroid differentiation. scaRNA tracks were scaled to the sample with the lowest number of millions of mapped reads and a biological triplicate merged for visualization. Intronic RNA tracks were normalized by reads per million and a biological triplicate merged for visualization. Differential count analysis were performed on raw unnormalized data using DEseq2

### Enhancers modify transcription initiation to regulate gene expression

Although our data suggest that changes in gene expression are predominantly driven by changes in initiation of transcription, others have proposed that enhancers regulate gene expression via changes in Pol II pausing (Chen et al. 2017; Ghavi-Helm et al. 2014; Liu et al. 2013; Schaukowitch et al. 2014). To help resolve this dichotomy, we investigated the role that enhancers play in regulating promoter proximal transcription by studying the well characterized enhancer clusters, which meet the current criteria of superenhancers and regulate the α- and β-globin genes (Bender et al. 2012; Gariglio, Bellard, and Chambon 1981; Lee et al. 2015; Sawado et al. 2003; Vernimmen 2014).

The mouse α-globin locus is regulated by five enhancers (R1-R4 and Rm) (Figure 4A). Two of the enhancers (R1 and R2) when deleted in combination (ΔR1R2) reduce nascent α-globin expression by ∼90% (Hay et al. 2016). The β-globin genes are regulated by six enhancers in the mouse (HS1-HS6) (Figure 4A). Deletion of two of these enhancers (HS2 and HS3) in combination (ΔHS2HS3) results in a ∼70% reduction in nascent β-globin expression (Bender et al. 2012). Previous work from our lab suggested that the human α-globin enhancers promote transcriptional initiation via recruitment of components of the pre-initiation complex and Pol II (Vernimmen et al. 2007). By contrast, the β-globin enhancers have been reported to facilitate release of paused polymerase (Bender et al. 2012; Gariglio, Bellard, and Chambon 1981; Hay et al. 2016; Sawado et al. 2003). Therefore, the α- and β-globin genes provide an ideal test bed to understand the mechanisms by which enhancers regulate the transcription cycles of their target genes.

**Figure 4.**
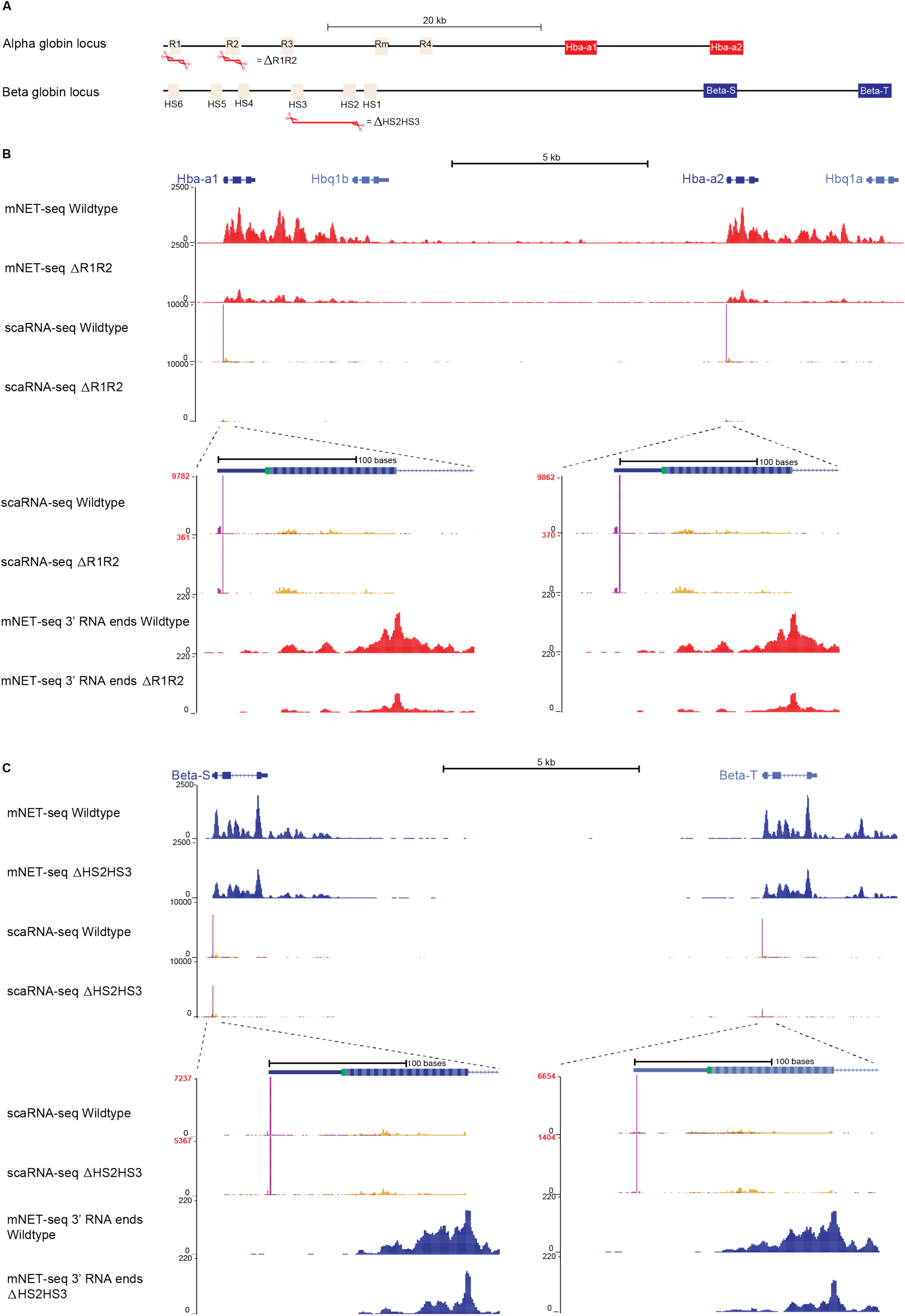
Enhancers regulate target gene expression via transcription initiation. (A) The α- and β-globin loci represented as a schematic and to scale. Regulatory elements (enhancers) are indicated as orange boxes and the two copies of α-globin at the α-locus are highlighted in red. Two copies of β-globin at the β-globin locus are highlighted in blue. At the α-globin locus the R1 and R2 enhancers when deleted in combination (ΔR1R2) reduce nascent α-globin expression by 95% (Hay et al. 2016). At the β-globin locus the HS2 and HS3 enhancers when deleted in combination (ΔHS2HS3) reduce nascent β-globin expression by 70% (Bender et al. 2012). (B) The α-globin locus in wildtype and ΔR1R2 cells, with the two α-globin genes (Hba-a1 and Hba-a2). scaRNA-seq and mNET-seq signals across the gene are both reduced showing that in the ΔR1R2 model there is a loss of transcription initiation. Two zoomed in views, highlight each genes promoter proximal region, with scaRNA-seq and mNET-seq. mNET-seq data represents the distribution of 3’ most base of each read (representing Pol II active site) with a 2nt window, to smooth data for visualization. scaRNA-seq data on zoomed view uses a scaled Y-axis (red values) to aid in visualization of the profile of Pol II pausing (3’ scaRNA ends). scaRNA-seq data on zoomed view uses a scaled Y-axis (red values) to aid in visualization of the profile of Pol II pausing (3’ scaRNA ends). (C) The β-globin locus in wildtype and ΔHS2HS3 cells, with the two β-globin genes (Beta-S and Beta-T). scaRNA-seq and mNET-seq signals across the gene are both reduced showing that in the ΔR1R2 model there is a loss of transcription initiation. Beta-T appears more affected than Beta-S. Two zoomed in views, highlight each genes promoter proximal region, with scaRNA-seq and mNET-seq. mNET-seq data represents the distribution 3’ most base of each read (representing Pol II active site) with a 2nt window, to smooth data for visualization. scaRNA-seq data on zoomed view uses a scaled Y-axis (red values) to aid in visualization of the profile of Pol II pausing (3’ scaRNA ends). scaRNA-seq and mNET-seq tracks were scaled to the sample with the lowest number of millions of mapped reads and a biological triplicate merged for visualization. mNET-seq data were normalized by merging a replicate (n=2 for each genotype) and subsampling to the data set with the lowest total read count and visualization.

To determine the effects of enhancer deletions on initiation and Pol II pausing, we performed scaRNA-seq in wildtype, ΔR1R2 homozygous mice (α-globin locus) and ΔHS2HS3 homozygous mice (β-globin locus) using primary erythroid cells derived from fetal liver. Deletion of enhancer elements at both loci results in a substantial decrease of scaRNA-seq transcripts at their TSSs (Figure 4B and 4C) indicating that in the absence of their enhancers, these genes either fail to initiate production of capped transcripts or that these transcripts are more rapidly turned over (increased abortive initiation). Equally affecting both α globin genes and predominantly the Beta-T gene at the β locus. In Wildtype, the globin genes in both loci exhibit an accumulation of RNA 3’ ends downstream of the promoters, indicative of Pol II pausing (Nechaev et al. 2010), yet increased levels of Pol II pausing does not occur in the enhancer knockouts, as this would have resulted in an increased level of paused/initiated scaRNA transcripts compared to nascent transcripts in the enhancer knockouts (see model in Figure 1B). To confirm and extend our previous observations that the ΔR1R2 deletions decrease the nascent output of the α-globin genes (Hay et al. 2016), we performed mNET-seq to measure RNA associated with Pol II through immunoprecipitation and sequencing (Nojima et al. 2015) to assess perturbation of the transcription cycle downstream of the promoter proximal region. These data corroborate our observations with scaRNA-seq. mNET-seq shows a uniform loss of both promoter proximal and elongating transcription. This shows that the enhancer deletions do not induce increased Pol II pausing but rather lead to a total reduction in RNA produced by Pol II at all points along the genes (Figure 4B and 4C).

### The mechanism by which enhancers regulate the initiation of transcription

We considered the potential mechanisms by which enhancers might affect the earliest stages of the transcription cycle. At many genes, close physical contact between the enhancer and promoter is associated with gene activation. We have previously shown that the mouse α-globin enhancers maintain contacts with their promoters when individual or pairs of enhancers are deleted from the cluster of enhancers using Next Generation Capture-C (Hay et al. 2016; Figure 5A). Here we show, that this is also the case at the mouse β-globin cluster (Figure 5B). It has also been shown using ATAC-seq and DNase assays respectively that the α- and β-globin enhancers are not required to maintain nucleosome free promoters. We confirmed these observations in both the ΔHS2HS3 (β-globin) model and the ΔR1R2 (α-globin) models (Hay et al. 2016; Figure 5A, B). Therefore, we conclude that the severe reduction in promoter proximal and full-length transcription seen in the absence of the globin enhancers is not due to a failure to form accessible chromatin or chromatin loops.

**Figure 5.**
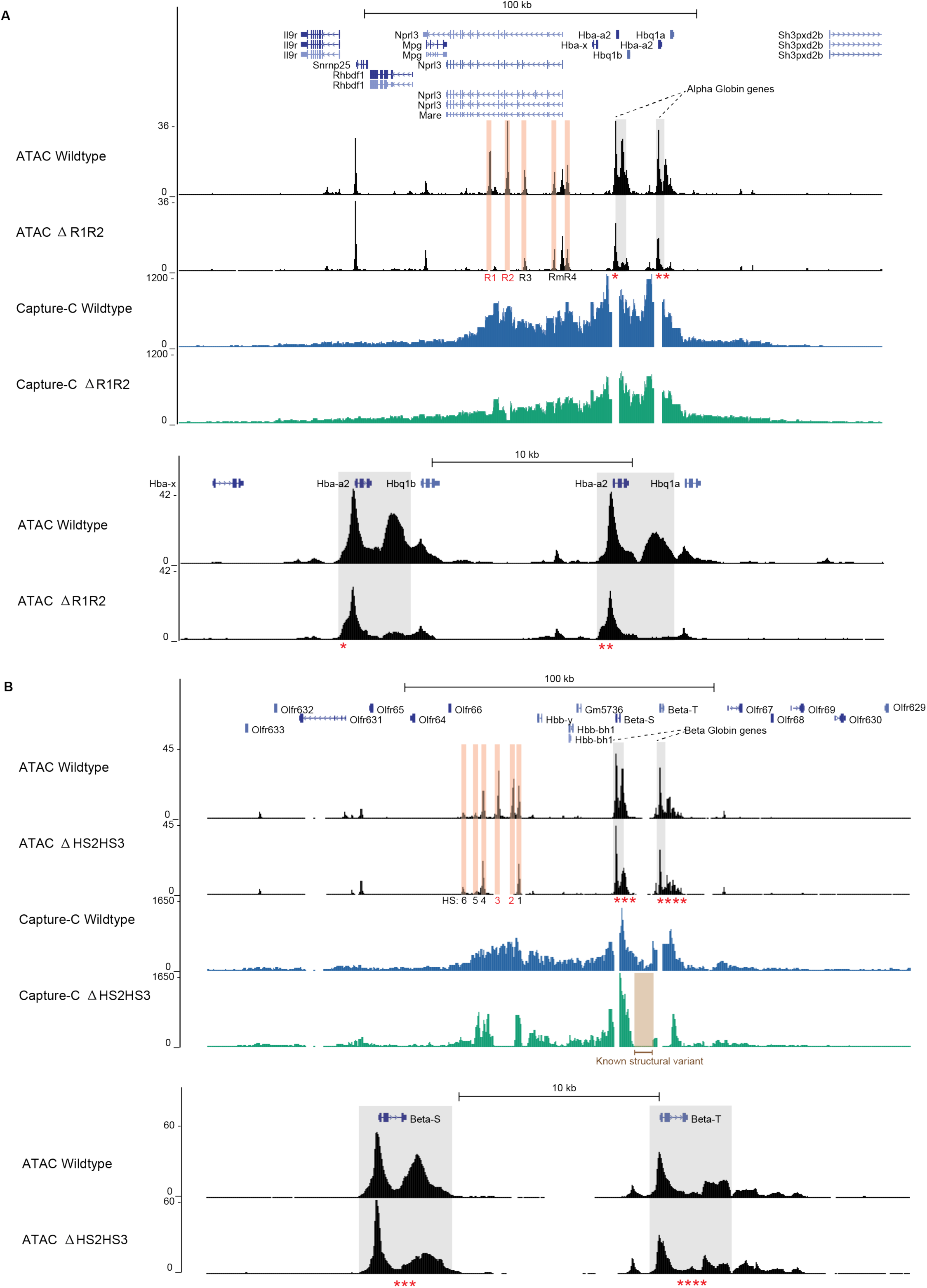
Promoter hypersensitivity and 3D chromatin structure are unperturbed by enhancer deletion. (A) α-globin locus with two copies of α-globin (Hba-a1 and Hba-a2 – grey boxes) and regulatory elements (R1-R4 and Rm) highlighted as orange boxes. Deleted enhancers (R1 and R2) are shown in red text and deletions are visible in the ATAC-seq data as lack of coverage when compared to wildtype. NG-Capture-C data show chromatin interactions from the viewpoint of the α-globin gene promoters (which appear as gaps in the Capture C track) interactions between the enhancers and the genes persist in the ΔR1R2 deletion suggesting chromatin hub formation or maintenance has not been affected by their deletion. Below, a zoomed in view of the α-genes shows that the promoters remain hypersensitive in the absence of the R1 and R2 enhancers. ATAC-seq signal in the 3’ UTR of the α-globin genes is also decreased in the ΔR1R2 mouse. (B) β-globin locus with two copies of β-globin genes (Beta-S and Beta-T marked as grey boxes) and regulatory elements (R1-R6) highlighted as orange boxes. Deleted enhancers (HS2 and HS3) are shown in red text and deletions are visible in the ATAC-seq data as lack of coverage when compared to wildtype. NG-Capture-C data show chromatin interactions from the viewpoint of the β-globin gene promoters (which appear as gaps in the Capture C track). Interactions between the enhancers and the genes persist in the ΔHS2HS3 deletion suggesting chromatin hub formation or maintenance has not been affected by their deletion. A known structural polymorphism in the locus specific to this mouse strain is shown as a brown box and results in no NG-Capture-C coverage for the ΔHS2HS3 over this region but does not affect interpretation of the genes or enhancers. Zoomed view of the genes shown highlighting that promoters remain hypersensitive in the absence of R2 and R3 enhancers. All ATAC-seq data performed in triplicate were merged and normalized by reads per million. NG-Capture-C was also performed in triplicate and tracks show the mean interaction profile. NG-Capture C and ΔR1R2 ATAC-seq data for the α-globin locus are from published work (Hay et al. 2016).

We next considered that the α- and β-globin enhancers may act to recruit or stabilize the PIC and/or Pol II at nucleosome free promoters. Previous studies at the human α-globin gene cluster showed a reduction of components of the PIC and Pol II in the absence of the enhancers (Vernimmen et al. 2007). To further test this hypothesis, we performed ChIP-seq of total Pol II in wildtype, ΔR1R2, and ΔHS2HS3 lines. This revealed that in the absence of the enhancers, at both the α- and β-globin loci there is a substantial reduction in the total level of Pol II both at the promoter and across the gene, but no evidence for a specific promoter proximal block to elongation (Figure 6A and Figure 6B). Again, this suggests that in the absence of enhancers there is a failure to recruit or stabilize the PIC leading to decreased initiation of transcription.

**Figure 6.**
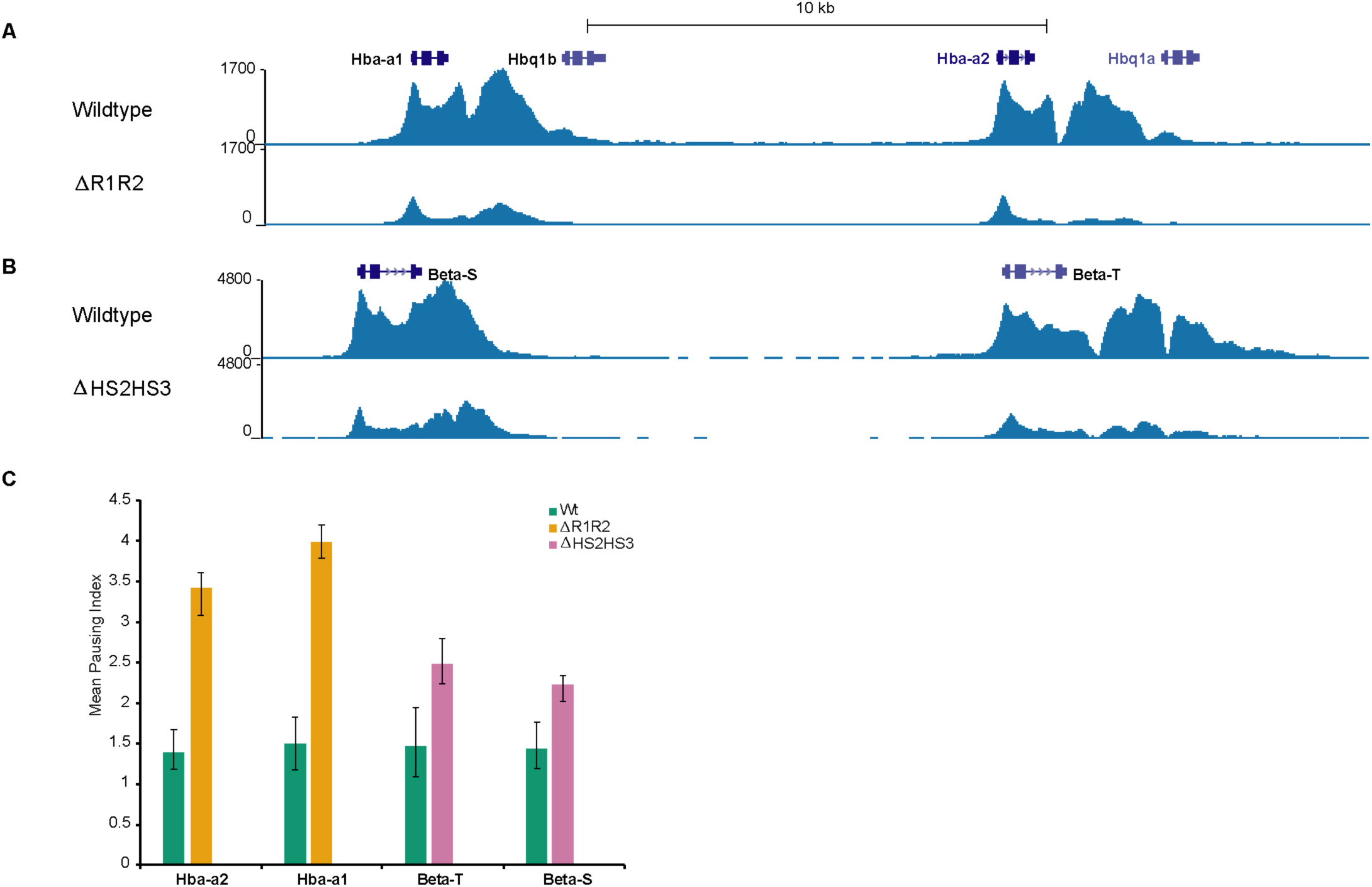
Enhancers stimulate Pol II recruitment to target genes. (A) Pol II ChIP-seq in wildtype and ΔR1R2 shows a total loss of Pol II across the α-globin genes (Hba-a1 and Hba-a2) consistent with defects in initiation not Pol II pause release) (B) Pol II ChIP-seq in Wildtype and ΔHS2HS3 shows a total loss of Pol II across the β-globin genes (Beta-S and Beta-T) consistent with a defect in initiation not Pol II pause release (C) Pausing index calculations (comparison of Pol II density per bp) at the promoter (TSS -100 to +300) versus the gene body (+301 to end of gene) showing α and β-globin have increased pausing indices, previously used to imply changes in Pol II pausing. All ChIP-seq experiments were performed in biological triplicate and each replicate normalized by millions of mapped reads under Pol II peaks before being merged for visualization. Pausing indices were calculated for each replicate and a mean (n=3) produced for each gene analyzed.

Previous studies of the ΔHS2HS3 mouse model used ChIP qPCR to study transcriptional pausing and elongation and concluded that the HS2 and HS3 enhancers function by increasing the transition from paused to elongating Pol II (Bender et al. 2012; Sawado et al. 2003). To investigate these differences in our conclusions we used the Pol II ChIP-seq data to calculate the pausing index of the individual copies of α- and β-globin genes in the wildtype and enhancer deletion lines (Figure 6C). The pausing index increases on average from ∼1.5 to ∼4 in the ΔR1R2 line and from ∼1.5 to ∼2.5 in the ΔHS2HS3 line, superficially indicating increased Pol II pausing (Rahl et al. 2010). Given the RNA-based evidence provided in this study showing a severe loss of both promoter proximal and gene body transcription, combined with the loss of Pol II recruitment at the globin promoters, we conclude that measurements of the Pol II pausing index derived from ChIP-seq alone do not reflect transcriptional pausing.

## DISCUSSION

Here we have shown that scaRNA-seq can simultaneously quantify transcriptional initiation and Pol II pausing and accurately identify TSSs and TPSs genome-wide. This has enabled us to re-interpret the process of RNA expression genome wide. It appears that the annotated TSSs for genes in current databases often lie upstream of the TSSs most commonly used in the widest number of cell types.

We show, using TSSs identified by scaRNA-seq rather than annotated TSSs, that most RNAs are initiated in a focused rather than dispersed manner and pause within a narrow window around +46bp downstream of the initiation site, juxtaposed to the +1 nucleosome (Figure 2C and E). This suggests that initiation and pausing are mechanistically related rather being independent regulatory steps in the transcription cycle. The tight coincidence between the accumulated paused RNAs and the stably positioned +1 nucleosome suggests it may form a significant transcriptional barrier.

The level of scaRNA-seq signal at any gene provides a direct quantitative assessment of promoter proximal transcription but on its own does not distinguish relative changes in initiation and pausing. However, in combination with a concomitant change in nascent transcription, the relative contributions of initiation and pausing become clear (Figure 1B). Using scaRNA-seq in combination with assays of nascent transcription we have shown that expression of RNA from most genes is regulated via the recruitment and/or initiation of transcription rather than via Pol II pausing. We further support this model of enhancer function using detailed studies of mouse models of the globin genes with and without their major enhancers and show that enhancers act primarily at the stage of recruitment and/or initiation.

The large uniform loses in RNA and Pol II at the globin genes in these deletion models, when contrasted with higher ChIP-seq based pausing indices, suggests that ChIP-seq derived pausing indices may increase due to lower levels of initiation, giving the false impression of higher levels of Pol II pausing (Figure 6).

This discrepancy may be explained by recent experiments, including live imaging *in vivo* which showed that most 5’ bound Pol II is rapidly turned over (Erickson et al. 2018; Kamieniarz-Gdula and Proudfoot 2019; Steurer et al. 2018) This could be associated with cycles of abortive initiation rather than stable pausing. Over time, such transiently associated Pol II may be detected by ChIP but would not necessarily produce scaRNAs which require transcriptionally engaged Pol II. This in turn could produce a distorted ratio of promoter proximal and elongating Pol II which does not reflect transcription *per se*.

In the context of an enhancer knockout, Pol II may still transiently associate with the promoter, albeit at much lower levels (Figure 6). As turnover of Pol II during recruitment/initiation is approximately two orders of magnitude faster than in gene bodies (J. Li et al. 2019; Steurer et al. 2018) this could result in a higher level of fixation in promoter versus gene bodies resulting in skewed increases in pausing indices which does not truly reflect the process of transcription which is readily seen using scaRNA-seq and nascent-RNA-seq.

Others have previously demonstrated the role pause release plays in gene expression by inhibiting factors such as NELF and DSIF. However, it is not clear that inhibitors of these factors are specific. DRB, one widely used inhibitor used to measure the extent of Pol II pausing in previous work, is in fact an ATP analogue that may inhibit other energetic processes in the cell, making conclusions from these studies difficult to interpret (Bensaude 2011). A further assumption is that the function of the protein targeted by chemical inhibition (CDK9) solely regulates Pol II pausing *in vivo*, disregarding off target effects on other processes in the cell which have been shown to include direct regulation of transcription termination through XRN2 (Sansó et al. 2016). Therefore, experiments using inhibitors may have over-estimated the role of Pol II pausing *in vivo* compared to our data, which comes from the differentiation of primary cells and *in vivo* mouse models without using such inhibitors. It is also reasonable to suggest that if CDK9 has roles beyond pausing, then inhibtion of this protein may have led to overestimates as to the prevalence and significance of Pol II pausing.

Interestingly, modeling of ChIP-seq density (Ehrensberger, Kelly, and Svejstrup 2013) has also suggested that promoter proximal peaks of Pol II need not be explained by a stable regulated pause but other kinetic mechanisms such as the dynamic interactions between recruitment, initiation and pausing of Pol II. This model also suggests that initiation must be saturated before Pol II can become regulated at the pause release step implying gene expression control using pausing would only be possible at extremely highly expressed genes.

In summary our data supports a model whereby levels of gene expression are primarily regulated via initiation and unexpectedly suggest that Pol II pausing as currently described, rarely contributes to differential gene expression. We suggest that rather than being an independent regulatory step, Pol II pausing represents an inherent kinetic bottleneck to transcription mediated by the +1 nucleosome, which results in premature termination of transcription. Additionally, our analysis of the function of the well-characterized α- and β-globin enhancers show that they act via the recruitment and/or transcriptional initiation of Poll II. Considering the lack of evidence, we find for regulation by pause release in these primary cells together this would suggest that enhancers in general control gene expression by regulating recruitment and/or initiation rather than promoter proximal pausing of Pol II.

## METHODS

### Data generation

#### Cell line culture

K562 cells obtained from the MRC WIMM transgenics facility were grown in RPMI media (ThermoFisher Scientific) supplemented with 15% FBS (37°C, 5% CO_2_). Cell density was maintained <=10^6^ cells per mL. Cell density and viability were determined using the NucleoCounter NC-3000 platform (Chemometec).

#### Mouse fetal liver culture

R1R2 deletion lines were maintained as heterozygotes and crossed to yield homozygous R1R2 deletions and wildtype litter mates as described in (Hay et al. 2016). HS2HS3 deletion lines carry a human YAC containing a rescue beta globin allele and are maintained as homozygotes as described in (Bender et al. 2012). Mouse fetal livers were harvested at E12.5 and disaggregated in expansion media (StemPro (ThermoFisher)) supplemented with Epo @ 1U/mL (Eprex, Janssen), SCF @ 50ng/mL (Peprotech), Dexamethosone @ 1μM (Hameln). Cells from individual fetal livers were grown for six days, maintaining a constant erythroblast cell count of <= 1 ×10^6^ cells per mL. Cells were genotyped as wildtype, homozygous R1R2 or homozygous HS2HS3 during expansion. Cells were counted using the NucleoCounter before selection for CD117 positive cells using magnetic assisted cell separation LS columns and CD117 MicroBeads following the manufacturers protocol (Milltenyi). Cells were left to recover for 6 hours in expansion media at a density of 1×10^6^ cells per mL. Aliquots of 5×10^6^ cells were harvested as the ‘0h’ timepoint corresponding to early erythropoiesis whilst the remaining cells were re-suspended in differentiation media (which is expansion media with the following changes: increased levels of Epo (5U/mL) and transferrin (0.5mg/mL), SCF and Dexamethasone removed). To induce erythroid differentiation cells were grown for 24 hours before harvesting for the ‘24h’ timepoint. At each time point, cell morphology was examined by preparing cytospins. Immunophenotypes were also determined by flourescence activated cell sorting using the Attune NxT (ThermoFisher Scientific) using anti Ter119 (PE-Cy7 labelled) and anti CD117 (PE labelled) antibodies (Biolegend). 0h of erythropoiesis was defined as (CD117+,Ter119-) and 24h as (CD117-, Ter119+). Isolated cells were confirmed as having these appropriate erythroid stage markers (Edling and Hallberg 2007; Hay et al. 2016; McGrath, Catherman, and Palis 2017).

#### Short capped RNA sequencing (scaRNA-seq)

5×10^6^ cells were aliquoted per replicate, pelleted by centrifugation (500g, 5min, R.T) and washed once in 5mL 1xPBS before lysis in TRI Reagent (Merck) by repeated pipetting. Samples were snap frozen in a dry ice ethanol bath and transferred to storage at -80°C until use. Samples were thawed on ice before being transferred to pre-spun (30s, 12,000g, R.T) phase lock gel tubes (VWR) and mixed with 1 volume of 1-bromo-3-chloropropane (Merck). Samples were spun (2min, 12,000g, R.T). The aqueous phase was transferred to a fresh 1.5mL Eppendorf tube and ethanol precipitated using 3 volumes of 100% ethanol, 0.1 volumes of 3M sodium acetate and 15μg GlycoBlue Coprecipitant (ThermoFisher Scientific). Samples were incubated (1hour, -80°C) before centrifugation (20,000g, 30min, 4°C). Pellets were washed with 1mL of ice cold 75% ethanol in DEPC treated water (ThermoFisher Scientific), and residual ethanol was removed by quick spin and pipetting. The pellets were resuspended in 13μL DEPC treated water and placed on ice. 0.5μL of RNA was taken and diluted 1:10 with DEPC treated water for QC. RNA integrity number (RIN) was determined using the RNA ScreenTape System (Agilent) according to the manufacturers protocol. RINs for all RNA samples used in this study were confirmed as being 9.6 or above (indicating negligible degradation of RNA prior to library prep). RNA species <=300nt were size selected on a 10% TBE-Urea gel (Thermo Fisher Scientific), which was prepared by pre-running for 15-20 mins at 180V. 4μL of the NEB ssRNA low molecular weight ladder was denatured in 36μL of NEB RNA loading dye for 4 mins at 90°C in a thermal cycler before snap cooling on ice. 12.5μL of extracted RNA in DEPC treated water was mixed with 12.5μL of Novex TBE-Urea Sample Buffer (2X) (Thermo Fisher Scientific) and denatured for 3 mins at 70°C before snap cooling on ice. The ladder and samples were loaded, skipping at least one lane between each sample. The gel was run for 50 mins at 180V (just before the lower size marker loading dye migrated to the bottom of the gel). The gel was then stained for 30 mins using SYBR gold (Thermo Fisher Scientific) in solution of 100mL 1x Ultrapure TBE (ThermoFisher Scientific). RNA 0-300nt in length was excised from the gel using scalpels and gel pieces placed into a 2mL Eppendorf tube with holes pierced in the bottom with a .22-gauge needle. The 2mL Eppendorf tube was then placed inside a 15mL falcon tube and the lid secured with parafilm. Samples were spun for (5mins, 4000g, 4°C) in a bench top centrifuge. Any remaining pieces of gel left in the Eppendorf were crushed through the pierced holes using a plunger from sterile 2mL syringe. The tube was inverted and flicked to dislodge and transfer any remaining gel pieces into the falcon. The falcon tube containing crushed gel was weighed and two volumes of elution buffer (Tris–HCl pH 7.5 20mM, NaAc 0.25M, EDTA pH 8.0 10mM, SDS w/v 0.25%) were added. The sample was vortexed and spun (1 min, 4000g, 4°C) in a bench top centrifuge. Samples were frozen for 15 mins at -80°C before being placed at R.T overnight to elute the RNA. 2mL of vacuum centrifuge grease (Corning) was added to a fresh 15mL falcon. The samples and fresh falcon were spun (5min, 4000g, R.T). The supernatant from the sample was transferred into the falcon with grease using a P1000 pipette with a wide bore pipette tip (Care was taken to minimize transfer of gel pieces to the fresh tube). A further 0.5 volumes of elution buffer was added to the sample gel pieces, before vortexing and spinning again (5 mins, 4000g, R.T). The second supernatant was transferred to the tube with the first. A phenol chloroform extraction was performed using the phase lock light tubes exactly as before except the final pellet for each sample was sequentially resuspended in total volume of 6μL DEPC treated water by pipetting. 3’ adapter ligation was performed for 1 hour and according the NEB small RNA kit for Illumina protocol. The reaction was topped up to 200μL with DEPC treated water and a phenol chloroform extraction performed using phase lock light tubes followed by and ethanol precipitation as before except the pellet was resuspended in 25.5μL DEPC treated water. A phosphatase treatment was performed using calf intestinal phosphatase (CIP, NEB) to remove fragmented RNA molecules from downstream library preparation for 30 mins @ 37°C. The reaction was topped up to 200μL with DEPC treated water and a phenol chloroform extraction performed as before except the pellet was resuspended in 26μL DEPC treated water. A decapping reaction was performed in a reaction volume of 30μL for 30 mins @ 37°C using CAP-clip (Tebu bio). The reaction was topped up to 200μL with DEPC treated water and a phenol chloroform extract performed (as before except the pellet was resuspended in 17.5μL DEPC treated water. The SR RT primer (NEB small RNA kit for Illumina) was hybridized to the 3’ adapter and the 5’ adapter ligated by adding 2μL T4 RNA ligase buffer (NEB), 5μL PEG 8000 (NEB), 1μL SR RT Primer and following the manufacturers protocol (NEB small RNA for Illumina library prep); “Hybridize the reverse transcription primer” followed by “ligate 5’ SR adaptor…”. A reverse transcription reaction was performed in a reaction volume of 50μL by adding Superscript III enzyme and buffers directly to the 5’ adapter ligated samples (in the proportions given in the SuperScriptIII protocol) and incubating for 1 hour at 50°C (following the manufacturers protocol). The reverse transcribed samples were subjected to PCR for 15 cycles by adding 50μL of the PCR mastermix from the NEB small RNA for Illumina library prep kit (NEB), 5μL of the Universal adapter and 5μL of a unique indexed adapter (see NEB library pooling guidelines for suitable adapter combinations for three sample sequencing runs). Library clean-up was performed using Ampure XP beads (Beckman) (using 110μL per PCR reaction and following manufacturer’s instructions). Libraries were eluted from beads into 23μL 1 x T.E (Tris-HCl pH 8.0 (10mM), EDTA pH 8.0 (1mM)).

K562 scaRNA-seq libraries were produced without replication for meta-analysis.

All mouse fetal liver scaRNA-seq libraries were produced in biological triplicate at each timepoint (wildtype 0h and 24h) and (HS2HS3 and R1R2 24h).

#### mNET-seq

mNET-seq was performed as in (Nojima et al. 2015) but with a staring input cell number of 5×10^6^. K562 mNET-seq libraries were produced without replication for meta-analysis.

All mouse fetal liver mNET-seq libraries were produced in biological duplicate at the 24h timepoint (wildtype HS2HS3 and R1R2).

#### RNA-seq

5×10^6^ cells were aliquoted per replicate, pelleted by centrifugation (500g, 5min, R.T) and washed once in 5mL 1xPBS before lysis in TRI Reagent (Merck) by repeated pipetting. Samples were snap frozen and transferred to storage at -80°C until use. Samples in TRI reagent were thawed on ice before extraction using the Direct-zol RNA Miniprep Kit (Zymo) according to the manufacturers protocol and including an on-column DNase digestion step for an increased incubation time of 30mins. The concentration of RNA in the sample was determined using the Qubit RNA BR Assay (ThermoFisher Scientific) and the RNA integrity number (RIN) was determined using the Tapestation and broad range RNA reagents. All RIN scores for samples used in this study were 9.6 or above (indicating negligible degradation of RNA prior to library prep). 2.0mg of Total RNA was depleted of ribosomal and globin RNA species with the Globin-Zero Gold rRNA Removal Kit (Illumina). Further fractionation of rRNA depleted RNA was performed using the NEBNext Poly(A) mRNA Magnetic Isolation Module (NEB) according to the manufacturers protocol (retaining the washes to use as input for polyA-library prep). The washes containing polyA-RNA are purified using isopropanol precipitation. ribosomal RNA depleted Total RNA, polyA+ RNA and polyA-RNA were then made into a strand specific library using the NEBNext Ultra II Directional RNA Library Prep Kit for Illumina (NEB) according to the manufacturers protocol (utilizing a 15-minute fragmentation step, 12 cycles of PCR using single indexing).

K562 scaRNA-seq libraries were produced without replication for meta-analysis.

Mouse fetal liver scaRNA-seq libraries were produced in biological triplicate at each timepoint (0h and 24h)

#### ChIP-seq

5×10^6^ cells were pelleted (500g, 5 mins, RT) before resuspension in 9mL of RPMI media (+10% FBS) and addition of 1mL fixation solution (50mM HEPES pH 8.0, 1mM EDTA pH 8.0, 0.5mM EGTA pH 8.0, 100mM NaCl, 10% formaldehyde). Samples were incubated on the roller at room temperature for 10 mins and fixation reactions quenched by the addition of 1.25mL of fresh glycine solution (1M). Samples were incubated on the roller for 5 mins before spinning down at (300g, 5 mins, 4°C), washing 1 x in 10mLs of ice-cold PBS and snap freezing the pellet in a dry ice ethanol bath. All samples were stored at -80°C until use. Cell pellets were removed from -80**°**C and lysed by the addition of 200μL cell lysis buffer (5mM PIPES pH 8.0, 85mM KCl, 0.5% IGEPAL-CA 630) and incubation on ice for 20 mins. Nuclei were pelleted (4**°**C, 900RCF, 10 mins), supernatant discarded and nuclei lysed in 100μL nuclear lysis buffer (50mM Tris-HCl pH 7.5, 10mM EDTA pH 8.0, 1% SDS) for 5 mins on ice. Chromatin was sonicated for 8 mins using a Covaris S220 sonicator (Duty cycle = 2%, Int = 3.0, Cycles/burst = 200, Power mode = frequency sweeping, Duration = 120s, Temp = 6**°**C). Insoluble material was pelleted by centrifugation (4**°**C, 20,000g, 20 mins), supernatant removed and 10x diluted with 1mL RIPA buffer without SDS (140mM NaCl, 10mM Tris-HCl pH 7.5, 1mM EDTA pH 8.0, 0.5mM EGTA pH 8.0, 1% Triton X 100, 0.1% Sodium deoxycholate). 100μL of the diluted chromatin was frozen to use as an “input” control. 110μL of a 1:1 Protein A and Protein G Dynabead slurry (ThermoFisher Scientific) was prepared by combining equal volumes of the Protein A and G beads and washing twice in low salt RIPA buffer with SDS (140mM NaCl, 10mM Tris-HCl pH 7.5, 1mM EDTA pH 8.0, 0.5mM EGTA pH 8.0, 1% Triton X 100, 0.1% Sodium deoxycholate, 0.1% SDS) using a magnetic Eppendorf rack to separate beads from supernatant. The remaining sonicated chromatin was pre-cleared 2x by incubation on a rotating platform (4**°**C, 10RPM, 1 hour) with 5μL of the A+G bead slurry. 10μL of Pol II antibody (N20 – sc899c, Santa Cruz) was conjugated to 100μL of the bead slurry by incubation on a rotating platform (4**°**C, 10RPM, 1 hour). The precleared chromatin and antibody conjugated beads were combined and incubated overnight on a rotating platform (4**°**C, 10RPM). In the afternoon of the following day (24 hour incubation) washing of the immunoprecipitate was performed using 0.5mL of the following buffers at 4**°**C on a magnetic Eppendorf rack; 2 x Low salt RIPA (Tris-HCl pH 7.5, EDTA pH 8.0 (1mM), EGTA pH 8.0 (0.5mM), Triton X-100 (1%), SDS (0.1%), Sodium Deoxycholate (0.1%), NaCl (140mM)), 2 x High salt RIPA (Tris-HCl pH 7.5, EDTA pH 8.0 (1mM), EGTA pH 8.0 (0.5mM), Triton X-100 (1%), SDS (0.1%), Sodium Deoxycholate (0.1%), NaCl (0.5M)), 1 x LiCl RIPA (Tris-HCl pH 7.5, EDTA pH 8.0 (1mM), EGTA pH 8.0 (0.5mM), Triton X-100 (1%), SDS (0.1%), Sodium Deoxycholate (0.1%), LiCl (250mM)), 2 x T.E (Tris-HCl pH 8.0 (10mM), EDTA pH 8.0 (1mM)). For each wash step; tubes were spun briefly at 100g for 2s to avoid beads sticking to lid, placed in magnetic rack and allowed to clear for ∼1min. Supernatant was removed and discarded, before removing tubes from the magnetic rack and adding 0.5mL of relevant buffer. Samples were incubated on the rotating platform @ 4°C for 4 mins. During the final wash incubation, a fresh solution of elution buffer (Tris-HCl (pH 7.5) (20mM), EDTA (pH 8.0) (5mM), NaCl (50mM), 1% SDS) was prepared for each immunoprecipitate and its corresponding input sample (0.6mL total per ChIP experiment). Sample supernatant was removed and beads resuspended in 0.15mL of elution buffer before transferring to a fresh Eppendorf tube. The input samples were thawed and 0.2mL of elution buffer added. The remaining elution buffer at room temperature until use the next day. To the beads and corresponding input 2µL or 1µL of RNAse A @ 10mg/mL (ThermoFisher Scientific) were added respectively. The beads and inputs were incubated on a thermomixer with shaking function (1,000rpm, 30s on, 30s off (interval mix)) for 30 mins, at 37°C. 1µL or 2µL of Proteinase K (20mg/mL) was added to the beads or inputs respectively and incubated on a thermomixer with shaking function (1,000rpm, 30s on, 30s off (interval mix)) at 65°C, overnight. Tubes were spun down briefly to collect droplets and transferred to a magnetic rack to separate. The supernatant was transferred to a new Eppendorf tube and beads were resuspended in 0.15mL of elution buffer before incubation for a further 5 mins at 65°C on the thermomixer. The quick spin and magnetic separation were repeated and the second supernatant transferred to the tube containing the first. A phenol chloroform extraction was performed using phase lock tubes followed by ethanol precipitation on samples and inputs. Samples and inputs were resuspended in 53μl 0.1 x T.E. Library preparation was performed using the NEB Ultra DNA library preparation kit for Illumina (NEB) and using 15 cycles of PCR followed by an Ampure XP bead (Beckman) clean-up.

Mouse fetal liver ChIP-seq libraries were produced in biological triplicate for all genotypes (Wildtype, aHom and bHom at the 24h timepoint).

#### ATAC-seq

ATAC-seq was performed exactly as in (Buenrostro et al. 2013) on 5 ×10^5^ cell aliquots.

All ATAC-seq experiments were performed in biological triplicate (Wildtype and HS2HS3 at the 24h timepoint).

#### NG-Capture-C

5 ×10^6^ HS2HS3 mouse fetal liver cells were fixed using a final concentration of 2% formaldehyde in PBS, spun down 500g 5 mins and placed into lysis buffer before freezing at -80°C. 3C libraries were prepared using *Dpn*II and standard methods as previously described (Davies et al. 2015) with the following modifications: no douncing was performed, all spins were performed at 300RCF, and after ligation intact nuclei were pelleted (15 min, 300RCF), supernatant was discarded, and nuclei were resuspended in 300 µL 1x T.E (Sigma) for phenol chloroform extraction. Digestion efficiency was determined by RT-qPCR with TaqMan and custom oligonucleotide, and ligation efficiency qualitatively determined by gel electrophoresis. Only 3C libraries with >70 % digestion efficiency was used. 3C libraries were sonicated to 200bp in a Covaris S220 and indexed with NEB Next Illumina library Prep reagents (NEB). Enrichment for specific viewpoints was performed with 70mer biotinylated oligonucleotides designed using CapSequm (http://apps.molbiol.ox.ac.uk/CaptureC/cgi-bin/CapSequm.cgi). Double capture was performed in multiplexed reactions with pools of oligonucleotides targeting either promoter proximal or promoter distal *DpnII* fragments with each oligonucleotide at a working concentration of 2.9nM.

NG Capture-C experiments were performed in biological triplicate (HS2HS3 at the 24h timepoint).

#### Library QC and sequencing

For all libraries, the modal DNA library size and library concentrations were determined using the D1000 reagents on a Tapestation (Agilent) and the Qubit dsDNA HS Assay Kit (Thermo Fisher Scientific) respectively. Libraries were quantified by qPCR using the NEBNext Library Quant Kit following the manufacturer’s instructions. Library concentration by qPCR was used to calculate dilution required to produce 6nM libraries for sequencing on the NextSeq500/550 High Output v2 kit (75 cycles) (Illumina). Sequencing was performed in paired end mode using 40 cycles of sequencing per read and a 6-cycle index read.

### Data analysis

#### Deriving annotated genes from UCSC table browser

Lists of Genbank, Ensembl, Refseq and UCSC genes were download from the UCSC table browser (http://genome.ucsc.edu/cgi-bin/hgTables) in .bed format for mm9 or hg19 genome builds.

#### Generation of unique UCSC gene and TSS lists

Gene isoforms were collapsed if their chromosome start and end coordinates matched using the script IdentifyUniqueUCSC.pl. This resulted in 60741 genes for further analysis. TSS coordinates were extracted using awk form the 5’ most coordinate of the genes in a strand specific manner.

#### Generation of non-coding RNA list

Lists of Ensembl non coding gene coordinates (for mm9 and hg19 genes respectively) were downloaded from the UCSC table browser (http://genome.ucsc.edu/cgi-bin/hgTables), using the following filtering terms; “miRNA”, “tRNA”, “snoRNA”, “Mt_tRNA”, “Mt_rRNA”, “rRNA”, “snoRNA”, “snRNA”. Lists of UCSC non-coding gene coordinates were also downloaded using the following filtering terms; “snRNA”, “miRNA”, “tRNA”, “snoRNA”, “Mt_tRNA”, “Mt_rRNA”, “rRNA”, “snoRNA”, “snRNA”. All coordinates were combined into a consensus list for either the mm9 or hg19 genome and windowed by 100nt up or downstream using bedtools slop.

#### Generation of splice sites list

A list of UCSC genes were downloaded from the table browser (http://genome.ucsc.edu/cgi-bin/hgTables) in gtf format and splice sites extracted from the exon coordinates using Awk.

#### scaRNA-seq

The script scaRNAseq_pipe.pl was used to perform the following analysis procedures: alignment of the data to the relative reference genome using Bowtie/1.1.2 (Langmead et al. 2009). This involved three alignment stages (first pass alignment, second pass adapter trimming of reads which did not map in first pass and realignment, third pass FLASH-ing (Magoč and Salzberg 2011) of reads which did not align second pass and realignment). All aligning reads were combined and remapped. Reads mapped in proper pairs were isolated from this file as a .bam file using samtools/0.1.19 view (H. Li et al. 2009) and flags (-bS -f 3). The bam file was converted into a .bedpe file using bedtools/2.25.0 (Quinlan and Hall 2010) bamtobed. Chromosome, start, end and strand of each read was inferred from the .bedpe file allowing RNA molecules to be “reconstructed” an output in .bed6 file using awk. Files were sorted (-k1,1 -k2,2n) and features overlapping non coding RNAs and annotated exon 3’ splice sites were removed using bedtools/2.25.0 intersect. Total read counts at this stage were used to subsample down to the sample with the lowest read coverage before (unix shuf). These files were converted into strand specific bigWigs of 5’ and 3’ most bases (four files in total) using bedtools genomecov and ucsctools bedGraphToBigWig (James Kent et al. 2002). bigWigs for the same genotype, strand and end are merged bigWigMerge (James Kent et al. 2002) before visualization on the UCSC genome browser.

For visualization of whole reconstructed RNA molecules, a region of interest was specified in .bed format and the script frags2bed.pl used to isolate RNA molecules overlapping this region entirely. RNA molecules were scored based on frequency to produce a color gradient before visualization on UCSC as .bed files. K562 scaRNA-seq data was not normalized prior to visualization.

scaRNA TSSs were called by creating a Homer Tag Directory (makeTagDirectory -format bed -sspe -keepAll -genome hg19/mm9) from a reconstructed RNA molecule file and calling peaks with (findPeaks -style tss) (Heinz et al. 2010). All peaks mapping to X, Y, M and random chromosomes were removed using grep -v.

Meta profiles of scaRNA-seq coverage of TSSs were produced using Homer/4.8 (Heinz et al. 2010) by converting the reconstructed RNA molecules file “into a “tag directory” makeTagDirectory.pl and the following flags (-format bed -sspe -keepAll -genome hg19). The distribution of 5’ or 3’ ends of reads, was plotted in 1bp bins around a given list of TSSs (+/-) 500bp) in a strand specific manner using annotatePeaks.pl and the following flags (-hist 1 -size 1000 -pc 3 -len 1 -raw).

Observed TSS were defined as those which were located +/-500bp from the 5’ end of an annotated unique UCSC gene using bedtools/2.25.0 closest. The output from this process was parsed with a custom Perl script (PlotObsVsAnn.pl) which produces a histogram plot of TSS per bp relative to the annotated TSS (+/- 500bp) “Observed vs Annotated TSS”. Annotated TSS were defined as the original 5’ end coordinate of the gene for which an Observed TSS could be identified.

Motif enrichment around observed or annotated TSSs was performed using a custom Perl script (SeqPile2.pl). The coordinates of the TSSs were used to retrieve sequence information for a user specified window around this position. The sequence was scanned in a user specified bin size (1bp) to identify specific user specified motifs (also using the reverse of the user specified motif to account for strand). The sum of the motif occurrence was plotted for each bin, using the 5’ end of the motif as the point plotted (i.e. a 2nt motif ends at the TSS are plotted at -2 relative to the TSS).

A consensus list of observed mouse fetal liver TSS coordinates was produced by calling TSS in each of the biological triplicates for the early (0h) and late (24h) erythroid stages (n=6 peaks calls) resulting in n= 65451 TSS. These TSSs were merged to remove overlapping calls with identical coordinates resulting in n = 52571 TSS using the script CollapseTSSsummitDups.pl. These TSSs were further processed to remove TSSs mapping to X, Y, M or random chromosomes resulting in n= 51161 TSS. A minimum coverage filter (7 RNA molecules per TSS) was set to remove low coverage TSSs, this was determined based on coverage required to detect a given TSS across at least 3 replicates and resulted in n = 12237 TSSs.

The closest UCSC annotated gene was identified using bedtools closest, showing that the vast majority of called TSS fell within a window of -100 to +300 relative to the annotated TSS (Figure S3D). Therefore, a window of -100 to +300 was drawn relative to each UCSC gene and genes which had multiple called TSS associated with them collapsed to form consensus list of TSS windows in which to calculate differential RNA molecules coverage (n= 4760). The coordinates of introns of the identified genes were extracted from annotated introns (UCSC table browser) and converted into. gtf format.

Focused vs dispersed TSS analysis was performed by calling TSS in K562 scaRNA-seq data (n=18331) and selecting those which were within +/-500bp of an annotated gene’s 5’ end using bedtools/2.25.0 closest. The genes annotated 5’ was used as the “annotated” TSS. For each TSS reconstructed RNA molecules were counted in the window +/-500nt relative to the TSS center. The number of reconstructed RNA molecules falling in the window +/-2nt relative to the TSS were also counted. If >=50% of the total molecules fell within +/-2nt of the TSS (observed or annotated) that TSS was classed as focused if <50% total reads fell in this region the TSS was classed as dispersed. The relative percentages of focused or dispersed TSS for annotated or observed TSS were plotted as a bar graph in Microsoft Excel.

Heatmaps of scaRNA-seq 5’ and 3’ ends were generated by producing bed files containing only the 5’ or 3’ most base of each reconstructed RNA molecule using awk. These files were then plotted with a custom R script (plot_heatmaps_tss_coverage.R)

Differential count analysis of mouse fetal liver scaRNA-seq was performed on counts of reconstructed RNA molecules which mapped to the window (−100 → +300 relative to a given annotated genes’ TSS) using bedtools/2.25.0 coverage and a custom script (deseq.html). TSSs with significantly differential counts (expression) were identified using a p value and false discovery threshold of <= 0.05. TSSs which had sufficient coverage to be analyzed were paired to a UCSC gene if one was located - 100+300 relative to the TSS.

#### NG-Capture-C

Data were analyzed using scripts available at https://github.com/Hughes-Genome-Group/CCseqBasicF/releases and R was used to normalize data (Davies et al. 2015).

#### ChIP-exo

Four replicates of Pol II ChIP-exo targeting Pol II (N20 antibody - sc899c) were downloaded from two separate studies; GSE108323 (Mchaourab, Perreault, and Venters 2018) and SRA067908 (Pugh and Venters 2016). The script ChIPexo_pipe.pl was used to align data to hg19 using bowtie/1.1.2 (Langmead et al. 2009), convert into .bam format and remove PCR duplicates using samtools/0.1.19(H. Li et al. 2009). The resultant .bam file was converted into a “tag directory” using Homer/4.8 makeTagDirectory.pl and the following flags (-format sam -sspe -keepAll -genome hg19). The distribution of 5’ ends of reads, was plotted in 1bp bins around a given list of TSSs (+/-) 500bp) in a strand specific manner using Homer/4.8 annotatePeaks.pl and the following flags (-hist 1 -size 1000 -pc 3 -raw). Distribution of data on the reverse strand was multiplied by -1 to flip the data on the X axis and allow easier visualization of regions occupied by Pol II.

#### RNA-seq

Published data were downloaded in .fastq format and reads trimmed to a maximum length of 50bp using awk to remove lower quality bases from 3’ ends of reads and improve alignment. Data generated in this study utilized 40bp paired end sequencing and so reads were not trimmed. The script alignment2Intronic.pl was used to align all data to the relevant reference genome (mm9 or hg19) using bowtie/1.1.2 (Langmead et al. 2009) and isolate reads which map 100% to an intron and do not map to an annotated non-coding RNA (see generation of non-coding RNA lists). The total number of reads in the resultant bam file was used to derive a reads per million (RPM) scaling factor which was applied when generating strand specific bedGraph coverage files with bedtools/2.25.0 (Quinlan and Hall 2010). bedGraphs were converted into bigWig format using ucsctools (Kent et al. 2010). bigWig files from duplicates were merged prior to visualization on the UCSC genome browser (James Kent et al. 2002).

polyA+ RNA-seq in K562 data were analyzed by using the script alignment2allreads.pl, wherein data alignment to the reference genome is performed with STAR/2.4.2a (Dobin et al. 2013), only reads which did not map to annotated non coding RNAs were retained and data were converted to bigWig without scaling by RPM.

Meta profiles of intronic reads coverage over refseq genes were generated from using ngsplots (Shen et al. 2014) and the following flags; -G hg19 -D refseq -R genebody -RB 0.05 -MQ 0 -SE 0 -L 5000 - VLN 0 -MW 2.5.

Correlation analysis of RNA-seq techniques over annotated introns (obtained from the UCSC genome browser) was performed by counting reads mapping to introns using featureCounts (Liao, Smyth, and Shi 2014) and the following flags (-M -s 2 -O --ignoreDup). Mean read per kilobase per million read (RPKM) values were calculated for each intron in each technique using Microsoft Excel. Mean RPKM values were exported as a .csv file and correlation analysis and plotting of data as a “hexbin” performed using the script (Hexbin.py).

Differential count analysis of mouse fetal liver intronic RNA-seq was performed by counting reads mapping to the introns of each annotated UCSC gene using featureCounts (Liao, Smyth, and Shi 2014) and the following flags (-M -s 2 -O --ignoreDup). Changes in counts were calculated using the script (deseq.htmL). Genes with significantly differential counts (expression) were identified using a p value and false discovery threshold of <= 0.05. A gene was paired to an observed (scaRNA-seq derived) TSS if one could be located in the window of -100 → +300 relative to that genes annotated gene TSS.

#### ChIP-seq

The script ChIP_pipe_PEproperpairs.pl was used to align data to mm9 using bowtie/1.1.2 (Langmead et al. 2009), convert into .bam format and remove PCR duplicates using samtools/0.1.19 (H. Li et al. 2009). The resultant .bam file was converted into a .bedpe file of a reconstructed molecules based on coordinates of 5’ ends of read 1 and read 2 respectively. Input ChIP-seq data was peak called using MACS2 callpeaks, ignoring duplicate reads (Zhang et al. 2008). Peaks were called from aligned Pol II ChIP-seq bam files for each sample at each timepoint using the input sample as a control for treatment. A consensus list of Pol II peaks between two conditions (e.g. 3 x Wildtype versus 3 x R1R2) in all samples was made by combining the individual sample peak calls, merging features closer than or equal to 50bp on the same strand (bedtools merge -d 50 -s). Input specific peaks were removed from this consensus (bedtools intersect). Total millions of mapped reads under each peak for each replicate in the comparison were counted using bedtools coverage. Each sample was converted into a bigWig file scaled based on millions of mapped reads under peaks/ total millions of mapped reads (reads under peaks per million mapped reads). Triplicates for each condition were combined to produce a merged bigWig file for visualization on the UCSC genome browser.

#### ATAC-seq

Published and unpublished ATAC-seq data were analyzed using the script (ATAC_pipe.pl). In brief reads were aligned, PCR duplicates removed, read pairs converted into reconstructed DNA fragments based on coordinates of 5’ ends of read 1 and read 2, adjustment for Tn5 transposase cutting (forward strand fragments shift +4, reverse strand shift -5bp). Resultant .bed filed were converted into a reads per million mapped reads normalized bigWig using bedtools/2.25.0 (Quinlan and Hall 2010) and ucsctools (James Kent et al. 2002). Triplicates for each condition were combined to produce a merged bigWig file for visualization on the UCSC genome browser.

#### mNET-seq

The script mNETseq_pipe_withDupXYMandSSremoval.pl was used to analyze mNET-seq data. Reads were aligned to mm9 using Bowtie/1.1.2 (Langmead et al. 2009). Reads mapping in proper pairs were isolated as a .bam file using samtools/0.1.19 view (H. Li et al. 2009) and flags (-bS -f 3). PCR duplicates were removed with samtools rmdup. Read pairs overlapping non coding RNAs and annotated exon 3’ splice sites were removed using bedtools/2.25.0 intersect (Quinlan and Hall 2010) samtools view. The bam file was converted into a .bedpe file using bedtools/2.25.0 bamtobed. Chromosome, start, end and strand of each read was inferred from the .bedpe file allowing RNA molecules to be “reconstructed” an output in .bed6 file. Total fragment counts were used to generate normalized strand specific coverage tracks, which were scaled to the sample with the lowest fragment count. A coverage track of the 3’ most end of each fragment, windowed by +/-2bp, was generated. Duplicates for each condition were combined to produce a merged bigWig file for visualization on the UCSC genome browser.

Meta profiles of mNET-seq coverage relative to observed K562 TSSs was produced by converting the fragments file into a “tag directory” using Homer/4.8 makeTagDirectory.pl (Heinz et al. 2010) and the following flags (-format bed -sspe -keepAll -genome hg19 -flip). The distribution of 5’ or 3’ ends of reads, was plotted in 1bp bins around a given list of TSSs (+/-) 500bp) in a strand specific manner using Homer/4.8 annotatePeaks.pl (Heinz et al. 2010) and the following flags (-hist 1 -size 1000 -pc 3 -len 1 -raw).

#### MNase-seq

Published MNase data were downloaded and analyzed using a script (MNasePipe.pl). Reads were aligned to the hg19 genome, PCR duplicates removed, DNA fragments reconstructed based on read1 and read 2 coordinates and output as a .bed. Homer was used to produce a meta profile of MNase-seq data by producing a tag directory (makeTagDirectory -format bed -keepAll -genome hg19) and generate a coverage meta profile relative to observed or annotated TSS lists using annotatePeaks (-hist 1 -size 1000 -pc 3 -raw). Heatmaps of MNase-seq coverage relative of observed TSS were produced using a custom R script (plot_heatmaps_tss_coverage_vs2_with_mnase.R).

#### GRO-seq

Published GRO-seq data were downloaded and analyzed using a script (GROseq_pipe.pl). Reads were aligned to the hg19 genome using Bowtie, PCR duplicates and reads mapping to non-coding RNAs removed. Homer was used to produce a meta profile of GRO-seq data by producing a tag directory (makeTagDirectory -format bed -keepAll -genome hg19 -flip) and plotting read coverage relative to observed TSS using annotatePeaks (-hist 1 -size 1000 -pc 3 -raw -len 1).

#### GRO-cap

Published GRO-cap data were downloaded and aligned to the hg19 genome using Bowtie/0.1.12, the aligned .sam file was converted into a tag directory using Homer/4.8 (makeTagDirectory -format sam -keepAll -genome hg19). Meta profiles of GRO-cap data over TSS were plotted using Homer/4.8 annotatePeaks (-hist 1 -size 1000 -pc 3 -raw -len 1), with a list of UCSC annotated or scaRNA-seq observed TSS for each gene.

#### CAGE

Published CAGE data was downloaded in aligned .bam format from Encode. .bam files were converted to .bed using bedtools bamtobed. The .bed file was converted into a tag directory using Homer/4.8 (makeTagDirectory -format bed -keepAll -genome hg19). Meta profiles of CAGE data over TSS were plotted using Homer/4.8 annotatePeaks (-hist 1 -size 1000 -pc 3 -raw -len 1) using a list of UCSC annotated or scaRNA-seq observed TSS for each gene.

#### Pausing index calculations

Calculation performed based on formula from (Rahl et al. 2010) using a custom Perl script Calculate_TR.pl. Pol II density, in reads per bp, is calculated in the windows (−30+300) and (+301 to end of gene) relative to the annotated (UCSC) TSS. Promoter density is divided by gene body density to derive a “pausing index” for genes where both values are above 0.

#### ChRO-seq

ChRO-seq libraries were generated and analyzed as in (Chu et al. 2018) from 5×10^6^ primary mouse fetal livers.

ChRO-seq experiments were performed in biological duplicate for each model.

## DATA AND SOFTWARE AVAILABILITY

### Published data used in this study

K562: GRO-seq (GSE60456), ChIP-exo Pol II (GSE108323 and SRA067908), rRNA depleted polyA+ RNA-seq (GSE86660), rRNA depleted total RNA-seq (GSE90231), rRNA depleted poly-RNA-seq (GSE90231), MNase-seq (GSE78984). GRO-cap (GSE60453), CAGE (ENCFF623BZZ).

Mouse fetal liver: ATAC-seq (GSE78800) and Capture C (GSE78803).

All data are deposited in Gene Expression Omnibus under accession number: GSE138359

All scripts used in this analysis are available at: https://github.com/martinlarke/Enhancers-predominantly-regulate-transcription-initiation-in-vivo

UCSC sessions of the data are available:

http://genome-euro.ucsc.edu/s/martin_larke/0h%20and%2024h%20Mouse%20FL

http://genome-euro.ucsc.edu/s/martin_larke/24h%20Mouse%20FL%20Enh%20Del%20and%20Wt

An interactive MIG session to interrogate scaRNA/intronicRNA correlations is available https://mig.molbiol.ox.ac.uk/mig/cgi-bin/mig.cgi?proj_id=4760&rm=mode_2

## AUTHOR CONTRIBUTIONS

Conceptualization- Jim Hughes, Doug Higgs, Martin Larke;

Methodology - M.L, J.H, T.N, D.D, J.T;

Formal Analysis - M.L, J.T, J.H, R.S, R.B, M.O;

Writing Original Draft - M.L, D.H and J.H.;

Writing Review & Editing - N.P, M.B, D.H, J.H, D.D, M.O, R.B;

Funding Acquisition – J.H and D.H

## ACKNOWLEDGEMENTS

We thank J. Sloane-Stanley and J. Sharpe of the WIMM Transgenics Facility for mouse breeding and fetal liver provision, K. Clark, S-A. Clark and C. Waugh of the WIMM Flow Cytometry Facility for assistance in cell sorting. We wish to acknowledge the Centre for Computational Biology, MRC Weatherall Institute of Molecular Medicine, Radcliffe Department of Medicine and University of Oxford for use of their services in this project.

This work was supported by the Medical Research Council grant (MC_UU_0009/4 to D.R.H. and MC_UU_009/15 to J.R.H.). A Wellcome Trust Strategic Award (106130/Z/14/Z to D.R.H and J.R.H.). An ERC Advanced Grant [339270] and Wellcome Trust Investigator Award [107928|Z|15|Z] to T.N. and N.J.P. A Stevenson Junior Research Fellowship at University College, Oxford (to A.M.O.). A Sir Henry Wellcome Postdoctoral Fellowship (209181/Z/17/Z to R.A.B.). M.A.B was supported by NIH grant R37 DK44746 to M. Groudine.

---

**Figure S1.**
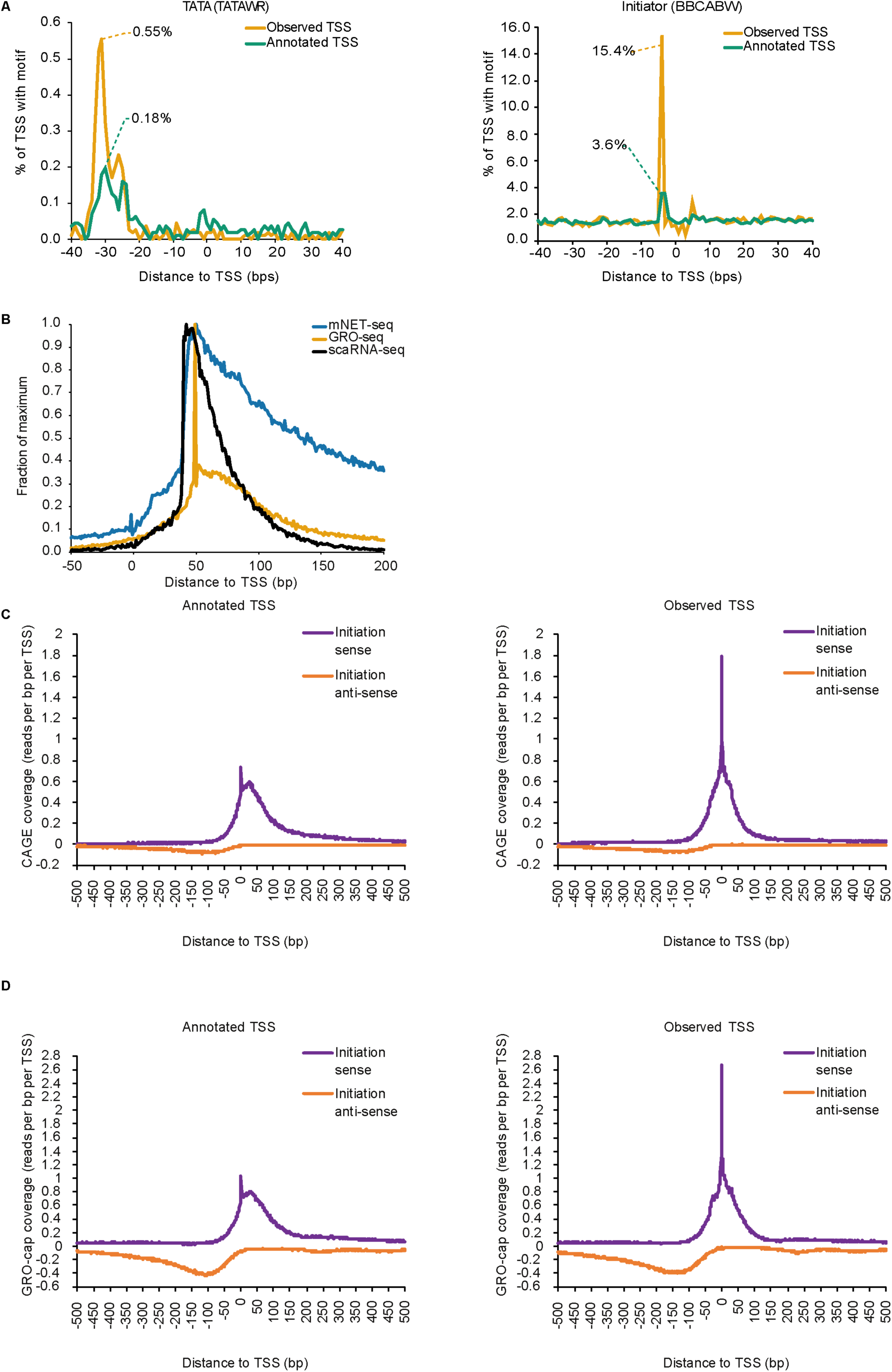
Further Validation of scaRNA-seq in mapping transcription initiation and pausing at a single molecule level *in vivo*. Related to Figure 1. (A) Enrichment of the TATA box (TATAWR) and Initiator (BBCABW) motifs (where B = C,G or T, W = A or T and R = A or G) in the window +/- 40bp relative to the scaRNA-seq observed (orange) TSSs or UCSC annotated (green) TSSs. The observed TSSs are called from scaRNA-seq data using Homer findPeaks in “tss” mode (Heinz et al. 2010). Such sites were retained if found within +/- 500bp of an annotated TSS associated with a UCSC gene (n= 11185 TSS). For comparison the TSS coordinates of these UCSC genes are used as the “annotated TSSs”. Data are from 5 × 10^6^ K562 cells. (B) A meta-analysis of 3’ RNA ends identified using scaRNA-seq, mNET-seq and GRO-seq relative to the Observed TSS (n= 11185 TSS). Only data mapping to the sense strand is displayed. All data-types produce similar profiles with maximum signal converging on 35-55nt downstream of the TSS (Pol II pausing). Data are plotted as a fraction of maximum coverage (reads per bp per TSS) for comparison. (C) A meta-analysis of CAGE data around annotated (UCSC) and observed (scaRNA-seq) TSSs, showing punctate initiation around the TSS and a loss of diffuse RNA 5’ ends signal downstream of the TSS after correcting for discrepancies in the positions of TSSs. (D) A meta-analysis of GRO-cap data around annotated (UCSC) and observed (scaRNA-seq) TSSs, showing punctate initiation around the TSS and a loss of diffuse RNA 5’ ends signal downstream of the TSS after correcting for discrepancies in the positions of TSSs.

**Figure S2.**
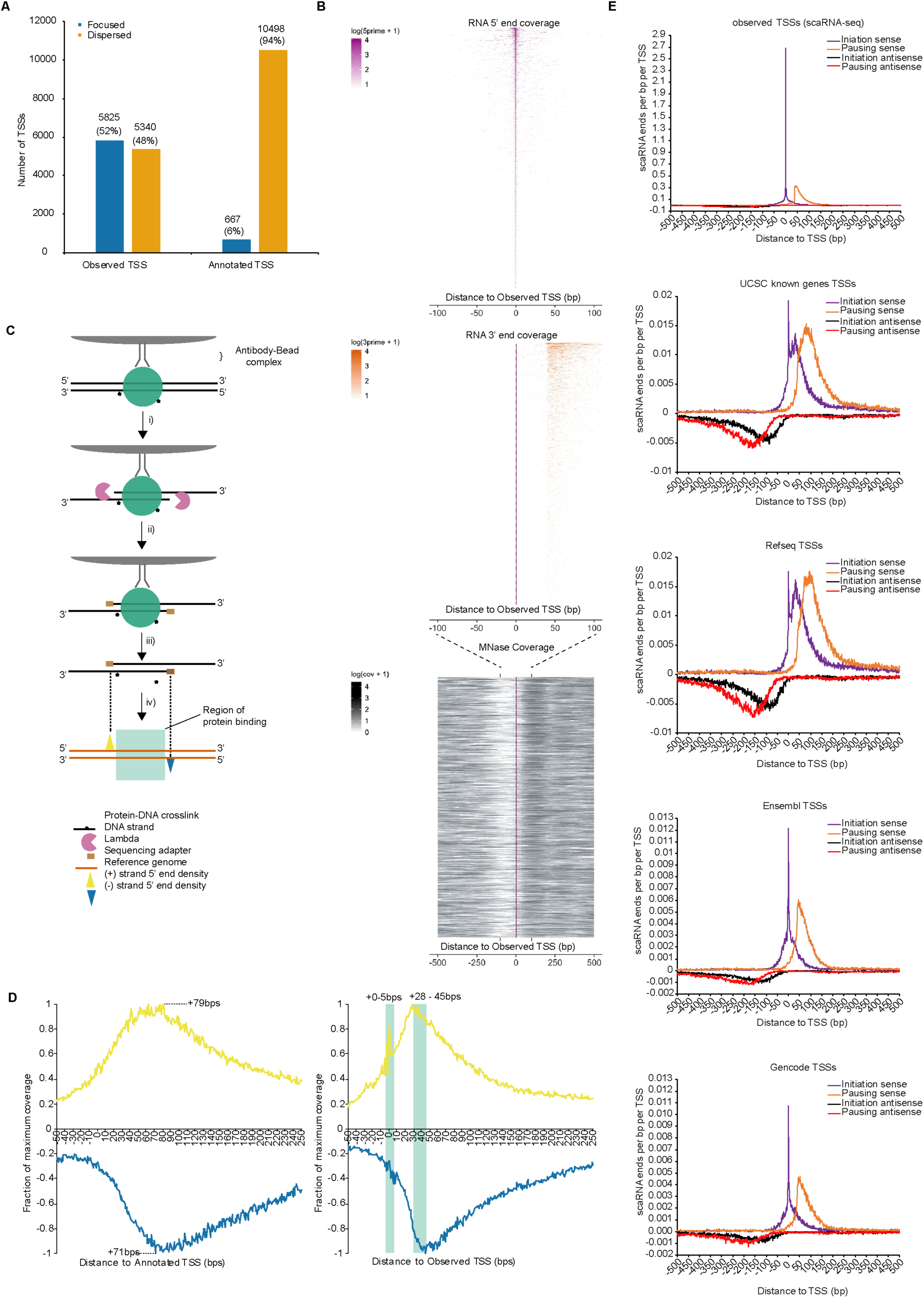
Correcting for the skew in scaRNA-seq observed TSSs compared to previously annotated UCSC TSSs revises the view of promoter proximal transcription. Related to Figure 2. (A) Analysis of the proportions of focused (>=50% of RNA 5’ ends in the window +/-2nt relative to the TSS) or dispersed (<50% of RNA 5’ ends in the window +/-2nt relative to the TSS) showing the use of annotated TSS coordinates results in the conclusion that there is widespread dispersed initiation. Correcting the annotated positions based on the scaRNA-seq observed positions results in approximately equal proportions of focused or dispersed TSSs. (B) Heatmaps of RNA 5’ ends (purple), RNA 3’ ends (orange). Coverage is per bp from the observed TSS on the sense strand. MNase-seq reads coverage over the TSS window +/-500 relative to the observed TSS position. Coverage is log (coverage +1) and capped at 55 reads per bp per TSS (log4) to aid in visualization of a wide dynamic range of coverage values as a heatmap. 0 bp position is highlighted on MNase and RNA 3’ end heatmaps to mark the observed TSS. (C) A schematic of the ChIP-exo protocol highlighting key stages. (i) on bead lambda exonuclease treatment of immunoprecipitated formaldehyde fixed chromatin, targeting a protein of interest such as Pol II. Ii) Ligation of a sequencing adapter to mark the point of exonuclease digest stop (boundary of the protein on DNA). Elution of DNA and library prep to retain directionality. (iv) Sequencing of library and mapping of reads, plotting only the 5’ most base (point of DNA-protein crosslinking) to resolve regions of protein binding. (D) Meta-analysis of Pol II Chip-exo data around observed or annotated TSS (n=11165) revealing that the observed TSSs produce a meta profile with more punctate peaks and evidence of Pol II occupancy at the TSS (−0-5bps) and Pol II pause site (+28-45bp). The use of annotated TSSs (left) produces an impossible combination of maximum signal at +79bps on the sense strand and +71bps on the antisense strand. The TSS peak is also not visible due to loss of resolution when using annotated TSSs. By contrast using the observed TSSs a pair of peaks on the sense and antisense strand between at +28 and +45bps respectively, delimits a region of Pol II occupancy (green). The same effect is also visible at the TSS (0-5bps) indicative of Pol II loading, a phenomenon only identifiable using the observed TSS. (E) A meta-analysis of scaRNA-seq data around the different TSS annotations. The skew evident in UCSC TSS annotation is also evident or more extreme in other commonly used annotations.

**Figure S3.**
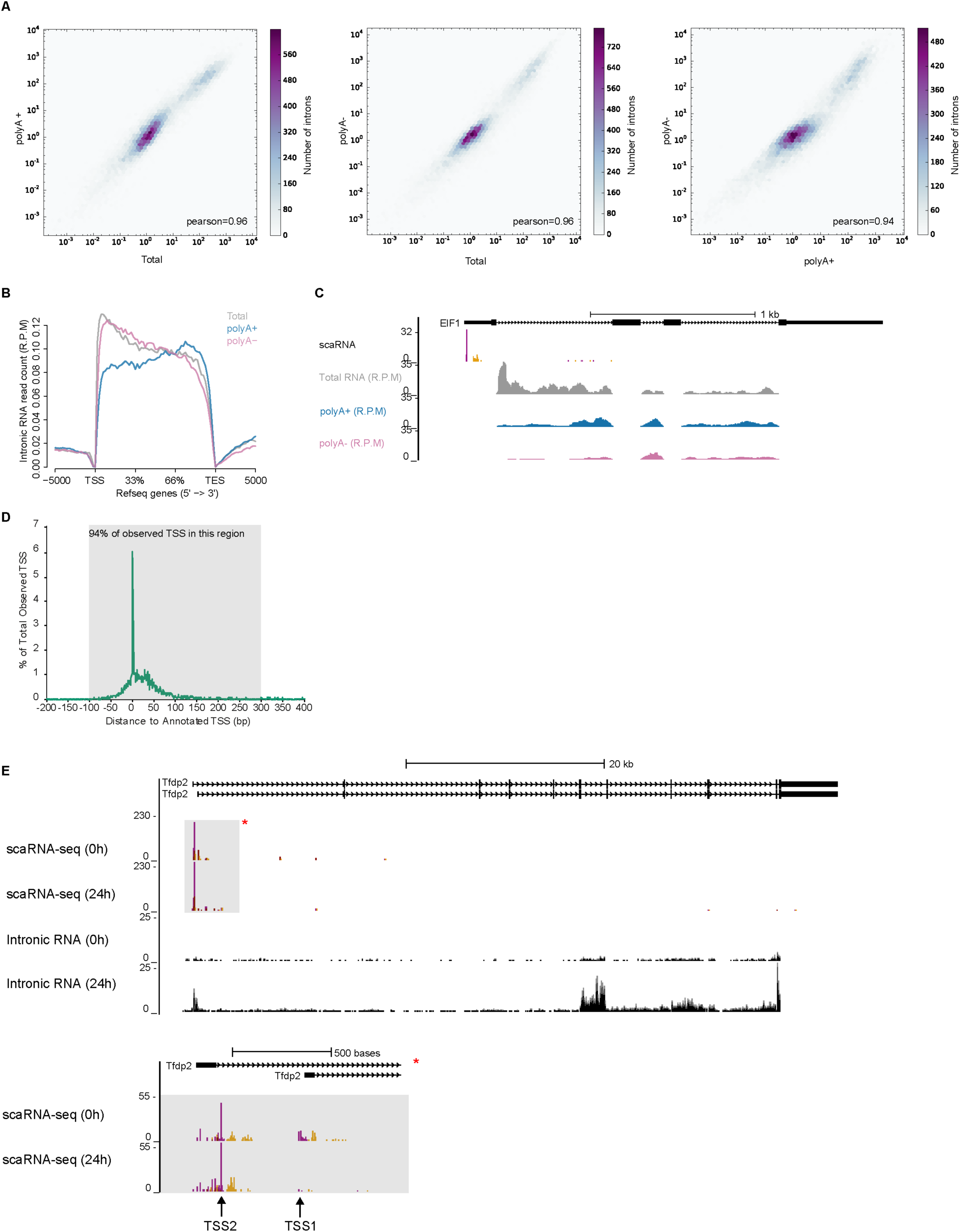
Intronic RNA as a measure of nascent transcription and windowing calculation. Related to Figure 3. (A) Correlation between intronic RNA coverage over all annotated introns (downloaded from UCSC table browser) comparing Total RNA, polyA+ and polyA-RNA data type and showing all data type correlate extremely highly (>= 0.94 pearson r^2^), suggesting they provide similar measures of intronic RNA. All coverage values are a mean of two isogenic duplicates. (B) Meta coverage of introns for each refseq gene. Data from total, polyA+ and polyA-types. Analysis shows that all data types provide comparable coverage over genes. With polyA+ showing the least 5’ coverage and most uniform coverage across genes. All coverage plots are a merge of two isogenic duplicates for each data types. (C) Example gene *Eif1* with scaRNA-seq, Total RNA (intronic reads only), polyA+ RNA (intronic reads only) and polyA- (intronic reads only) showing the most uniform coverage is generated from polyA+ RNA. (D) A plot of the distances between observed and annotated TSS, indicating that in primary mouse fetal liver genes observed TSS are systematically found downstream of their annotated positions (as in human (K562 cells). 94% of TSS are found in a window of - 100→+300 relative to the annotated TSS, defining these regions as suitable for counting scaRNA molecules for differential count analysis. (E) *Tfdp2* exemplifies increases in intronic RNA and decreases in scaRNA at 0h and 24h across the region of promoter proximal transcription (putative Pol II pausing). Close inspection of the promoter proximal region (grey box marked with red asterisk) reveals a loss of scaRNA signal at the TSS in question (TSS1) but a gain of signal at an alternative upstream TSS (TSS2), arguing against regulation of Pol II pausing.

